# CAMAP: Artificial neural networks unveil the role of codon arrangement in modulating MHC-I peptides presentation

**DOI:** 10.1101/2020.06.03.078824

**Authors:** Tariq Daouda, Maude Dumont-Lagacé, Albert Feghaly, Yahya Benslimane, Rébecca Panes, Mathieu Courcelles, Mohamed Benhammadi, Lea Harrington, Pierre Thibault, François Major, Yoshua Bengio, Étienne Gagnon, Sébastien Lemieux, Claude Perreault

## Abstract

MHC-I associated peptides (MAPs) play a central role in the elimination of virus-infected and neoplastic cells by CD8 T cells. However, accurately predicting the MAP repertoire remains difficult, because only a fraction of the transcriptome generates MAPs. In this study, we investigated whether codon arrangement (usage and placement) regulates MAP biogenesis. We developed an artificial neural network called Codon Arrangement MAP Predictor (CAMAP), predicting MAP presentation solely from mRNA sequences flanking the MAP-coding codons (MCCs), while excluding the MCC *per se*. CAMAP predictions were significantly more accurate when using original codon sequences than shuffled codon sequences which reflect amino acid usage. Furthermore, predictions were independent of mRNA expression and MAP binding affinity to MHC-I molecules and applied to several cell types and species. Combining MAP ligand scores, transcript expression level and CAMAP scores was particularly useful to increaser MAP prediction accuracy. Using an *in vitro* assay, we showed that varying the synonymous codons in the regions flanking the MCCs (without changing the amino acid sequence) resulted in significant modulation of MAP presentation at the cell surface. Taken together, our results demonstrate the role of codon arrangement in the regulation of MAP presentation and support integration of both translational and post-translational events in predictive algorithms to ameliorate modeling of the immunopeptidome.

**Author summary:** MHC-I associated peptides (MAPs) are small fragments of intracellular proteins presented at the surface of cells and used by the immune system to detect and eliminate cancerous or virus-infected cells. While it is theoretically possible to predict which portions of the intracellular proteins will be naturally processed by the cells to ultimately reach the surface, current methodologies have prohibitively high false discovery rates. Here we introduce an artificial neural network called Codon Arrangement MAP Predictor (CAMAP) which integrates information from mRNA-to-protein translation to other factors regulating MAP biogenesis (e.g. MAP ligand score and transcript expression levels) to improve MAP prediction accuracy. While most MAP predictive approaches focus on MAP sequences per se, CAMAP’s novelty is to analyze the MAP-flanking mRNA sequences, thereby providing completely independent information for MAP prediction. We show on several datasets that the integration of CAMAP scores with other known factors involved in MAP presentation (i.e. MAP ligand score and mRNA expression) significantly improves MAP prediction accuracy, and further validate CAMAP learned features using an *in-vitro* assay. These findings may have major implications for the design of vaccines against cancers and viruses, and in times of pandemics could accelerate the identification of relevant MAPs of viral origins.

## Introduction

In jawed vertebrates, virtually all nucleated cells present at their surface major histocompatibility complex class-I (MHC-I) associated peptides (MAPs), collectively referred to as the immunopeptidome [1,2]. MAPs play a central role in shaping the adaptive immune system, as they orchestrate the development, survival and activation of CD8 T cells [3]. Moreover, recognition of abnormal MAPs is essential to the elimination of virus-infected and neoplastic cells [4]. Therefore, systems-level understanding of MAP biogenesis and molecular composition remains a central issue in immunobiology [5,6].

The generation of the immunopeptidome can be conceptualized in two main events: (a) the generation of MAP candidates (i.e. peptides of appropriate length for MHC-I presentation) through protein degradation, and (b) a subsequent filtering step through the binding of MAP candidates to the available MHC-I molecules. Rules that regulate the second event have been well characterized using artificial neural networks (ANN) and weighted matrix approaches [7,8]. However, accurately predicting which peptides will ultimately reach MHC-I molecules following a multistep processing in the cytosol and endoplasmic reticulum remains an open question [6]. Most efforts at modeling MAP generation have focused on post-translational events and their regulation by the amino acid sequence of MAPs and of directly adjacent residues (typically 10-mers at the N- and C-termini). While the consideration of preferential sites of proteasome cleavage has proven useful to enrich for MAP candidates [9], it remains insufficient for MAP prediction, due to prohibitive false discovery rates [10–12].

A large body of evidence suggests that a substantial portion of MAPs are produced co-translationally [13–15], deriving from defective ribosomal products (DRiPs), that is, polypeptides that fail to achieve a stable conformation during translation and are consequently rapidly degraded.

This concept was initially supported by two observations: (i) viral MAPs can be detected within minutes after viral infection, much earlier than their associated proteins half-life [16], and (ii) MAP presentation correlates more closely with translation rate than with overall protein abundance [17,18]. In addition, while all proteins contain peptides that are predicted to bind MHC-I molecules, mass spectrometry analyses have revealed that the immunopeptidome is not a random excerpt of the transcriptome or the proteome [1,19]. Indeed, proteogenomic analyses of 25,270 MAPs isolated from B lymphocytes of 18 individuals showed that 41% of expressed protein-coding genes generated no MAPs [19]. These authors also provided compelling evidence that the presentation of MAPs cannot be explained solely by their affinity to MHC-I alleles and their transcript expression levels, while ruling out low mass spectrometry sensitivity as an explanation for the non-presentation of the strong binders. Because (i) MAPs appear to preferentially derive from DRiPs and (ii) codon usage influences both precision and efficiency of protein synthesis [20,21], we hypothesized that codon usage in the vicinity of MAP-coding codons (MCCs) might significantly contribute to the regulation of MAP biogenesis. We developed an artificial neural network called Codon Arrangement MAP Predictor (CAMAP), trained to identify MCCs flanking regions. We then used CAMAP to uncover key codon features that characterize mRNA sequences encoding for MAPs (i.e. source) when compared to sequences that do not (i.e. non-source).

## Results

### Dataset description

We analyzed a previously published dataset consisting of MAPs presented on B lymphoblastoid cell line (B-LCL) by a total of 33 MHC-I alleles from 18 subjects [19,22]. Because we were searching for features that influence MAP generation and not the binding of MAP to MHC-I molecules, we elected to analyze the MCC flanking sequences only and excluded the MCCs *per se* from our positive (hits) and negative (decoys) sequences (Fig. 1A). To facilitate data analysis and interpretation, we restricted our hit dataset to MAPs with a length of 9 amino acids, for a total of 19,656 9-mer MAPs (which represents 78% of MAPs in this dataset). We next created a decoy dataset from transcripts that generated no MAPs, by randomly selecting 98,290 9-mers from these transcripts. Finally, we used pyGeno [23] to extract MCCs flanking regions corresponding to both hit and decoy MAPs, which constituted our final dataset for CAMAP. Of note, each sequence in the final dataset is unique and derives from the canonical reading frame. In addition, in order to investigate the relative importance of codon vs. amino acid usage in MAP biogenesis, we generated a dataset of shuffled sequences (for both positive and negative datasets) in which original codon sequences were randomly replaced by synonymous codons according to their usage frequency in the dataset (Fig. 1B). This transformation was performed to ensure that both neural networks received the same number of parameters as input, preventing the introduction of a favorable bias for the codon network. The random shuffling causes any codon-specific feature to be shared among synonyms, thereby causing the shuffled codon distribution to reflect the amino acid usage (see Materials and Methods for more details). Indeed, codon distributions in the shuffled datasets more closely reflected those of their corresponding amino acid than in the original dataset (Supplementary Figure S1), with 92% of codons in the shuffled dataset showing a strong correlating (R^2^ > 0.95) with the amino acid distribution, compared to only 69% in the original dataset (p < 2×10^−16^, Supplementary Figure S2). Importantly, this shuffling does not affect the resulting amino acid sequence thereby preserving all potential amino acid-related motifs. Distributions of each codons in the original VS shuffled dataset and compared to its corresponding amino acid can be found in Supplementary Figure S3.

**Figure 1.**
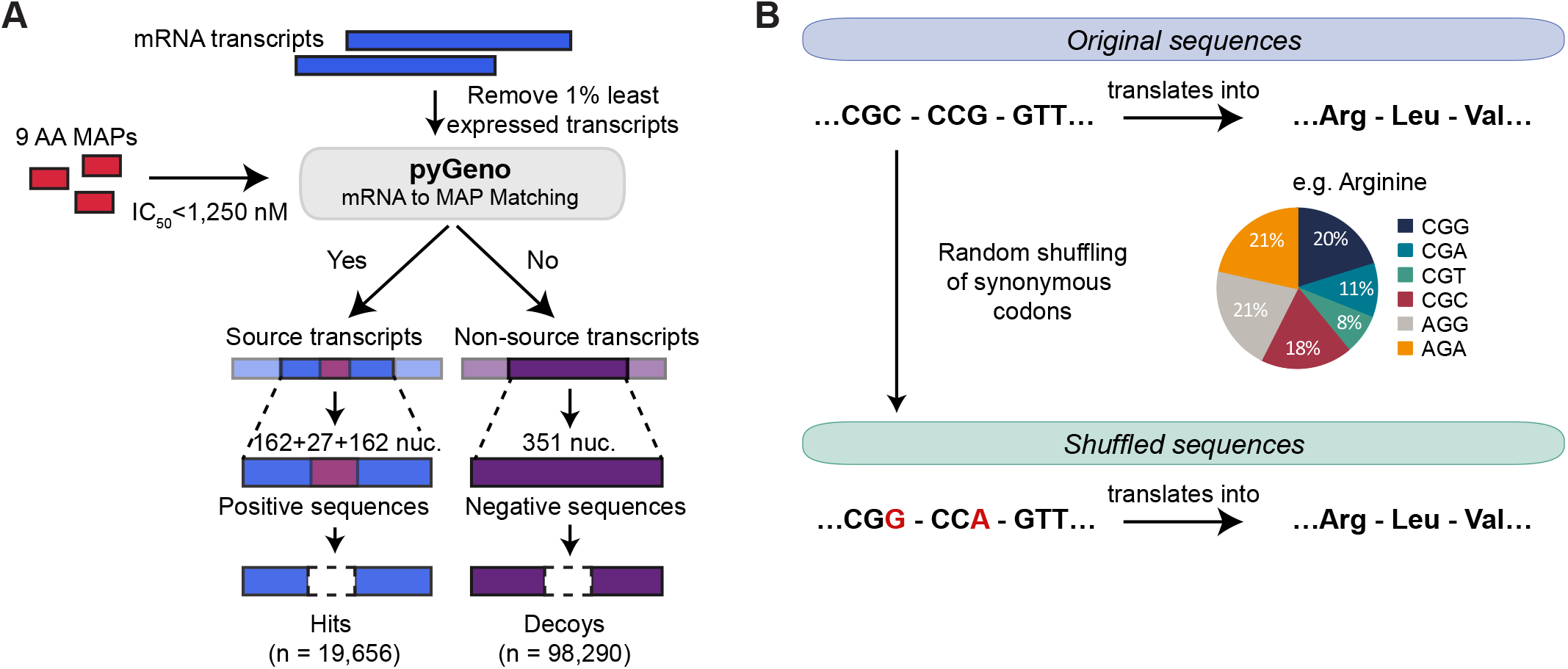
Construction of the dataset. (a) Transcripts expressed in B cells from 18 subjects were considered as source or non-source transcripts depending on their match with at least one MAP. Because we were searching for features that might influence MAP generation and not the binding of MAP to MHC-I, we focused our attention on mRNA sequences adjacent to the nine MCCs (i.e. up to 162 nucleotides on each side of MCCs). (b) Creation of the shuffled dataset. Codons were randomly replaced by a synonymous codon according to their respective frequencies (i.e. codon usage) in the dataset. The random shuffling causes any codon-specific feature to be shared among synonyms, thereby causing the shuffled codon distribution to reflect the amino acid usage. Importantly, both the original sequence and its shuffled version translates into the same amino acids.

### CAMAP links codon usage to MAP presentation

To assess the importance of codon usage in MAP biogenesis, we reasoned that if codons bear important information that is operative at the translational rather than the post-translational level, then: (i) CAMAP trained to identify MCCs flanking regions should consistently perform better when trained on original codon sequences than on shuffled codon sequences (reflecting amino acid sequences), and (ii) synonymous codons should have different effects on the prediction. To test these hypotheses, CAMAP received as inputs MCCs flanking regions from hit and decoy sequences from either the original or shuffled datasets. It was then trained to predict the probability that individual input sequences were MCCs flanking regions (i.e. hit) rather than sequences from the negative dataset (Supplementary Figure S4A).

We compared CAMAP performance when predicting MAP presentation from original codon sequences, versus shuffled sequences representing amino acid arrangement. To evaluate the robustness of our approach, 12 different CAMAPs were trained in parallel, with different train-validation-test splits of the dataset. Our results show that predictions were consistently better when CAMAP received the original codons rather than the shuffled sequences (Fig. 2A). CAMAPs receiving information from both pre-MCCs and post-MCCs sequences (i.e. whole MCC flanking context) also performed better than when receiving only pre- or post-MCCs context (Fig. 2A and Supplementary Figure S4B-C), suggesting that pre- and post-MCCs context are not redundant. Indeed, we found a weak correlation between the prediction scores of CAMAPs trained only with pre- or post-MCCs sequences (Supplementary Fig. S5). In addition, CAMAPs receiving longer sequences performed better than those receiving shorter sequences (Fig. 2B). Because sequences located far upstream and downstream of the MCCs (i.e. in ranges exceeding the direct influence of proteases) are informative regarding MAP presentation, it supports the existence of factors unrelated to protein degradation modulating MAP presentation.

**Figure 2.**
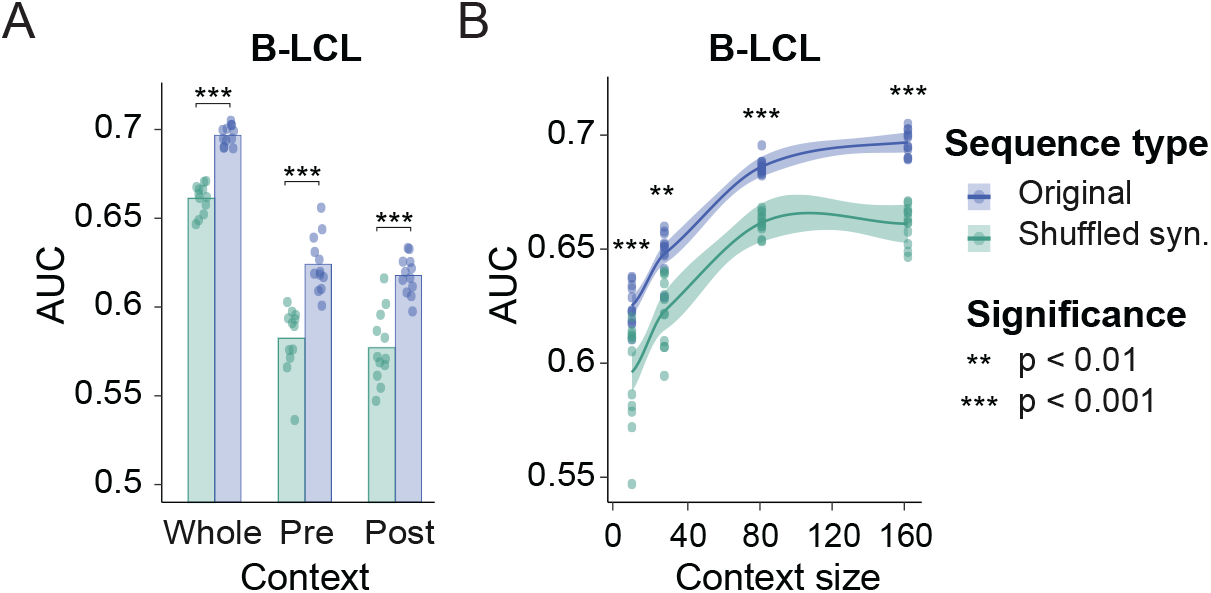
CAMAP predictions on MAP-flanking sequences. (A) Area under the curve (AUC) score for CAMAPs trained with whole MCCs context, versus CAMAPs trained with only pre- or post-MCCs context. All CAMAPs presented here were trained with a context size of 162 nucleotides. (B) AUC for CAMAPs trained with codon context sizes of 9, 27, 81 and 162 nucleotides (context here refer to mRNA sequences flanking the MCCs).

Both MAP binding affinity to the MHC-I molecule and the level of gene expression are predictive of MAP presentation [19]. Because codon usage has been shown to be different in highly expressed genes, we wanted to verify whether the codon-specific rules captured by CAMAP were associated with potential biases in our positive dataset, which is enriched in highly expressed genes. We first show that there is no correlation between gene expression levels and CAMAP scores in both the positive and negative datasets (R < 0.1, Fig. 3A). This was true for both average expression levels across our samples (Fig. 3A), and for samples individually (see Supplementary Fig. S6). Secondly, we trained CAMAP networks using a decoy dataset that mirrored the positive dataset gene expression level (Supplementary Fig. S7A) and showed similar results: CAMAP trained on original codon sequence performed better than CAMAP trained on shuffled sequences (Supplementary Fig. S7B). These results show that the codon-specific rules captured by CAMAP trained on original sequences are independent of gene expression levels.

**Figure 3.**
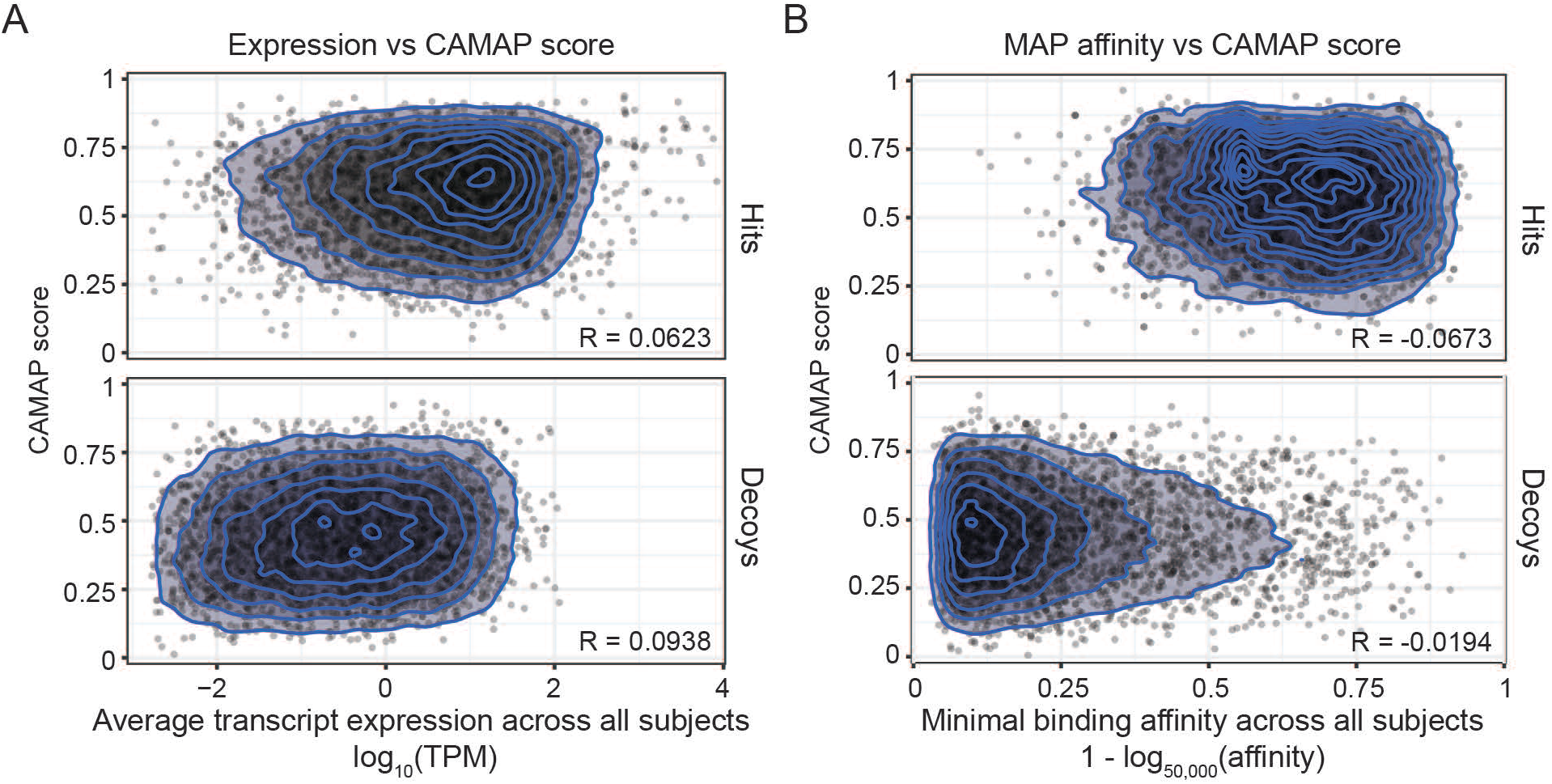
Correlation between CAMAP prediction score and (A) transcript expression levels and (B) MAP binding affinity. CAMAP used here was trained on original codon sequences using a context size of 162 nucleotides (both pre- and post-MCCs context).

We stipulate that the presence of MHC-I binding motifs in the MCCs in the positive dataset might be associated with biases in the MAP-flanking regions, which could also influence CAMAP training. Therefore, to evaluate the presence of this potential bias, we first evaluated the correlation between CAMAP scores and MAPs binding affinity. Again, our result showed no correlation between CAMAP scores and MAP binding affinity, both when considering the minimal binding affinity of each MAP to the MHC-I alleles contained in our dataset (Fig. 3B) or when considering each allele individually (Supplementary Fig. S8). Secondly, we trained CAMAP networks using a decoy dataset that mirrored the positive dataset MAP binding affinities (Supplementary Fig. S9A). Again, CAMAPs trained on original codon sequence performed better than CAMAPs trained on shuffled sequences (Supplementary Fig. S9B). These results show that codon-specific rules captured by CAMAP trained on original sequences are independent of MAP binding affinities and of potential biases in codon usage of MAP-flanking sequences associated with the presence of an MHC-I binding motif in the MCCs.

We next evaluated the possibility of biases associated with many MAPs originating from conserved regions (e.g., found in multiple domains of the same domain family such as zinc fingers or kinases). We first evaluated MAPs that could originate from different transcripts within the transcriptome (i.e. transcripts with sufficient expression levels detected by RNA sequencing) as they are likely to represent conserved regions in the genome. While 79.9% of MAP originated from unique contexts (Supplementary Fig. S10A), 2.1% of MAPs had more than 3 possible origins, which represented 11.7% of the hit dataset (Supplementary Fig. S10B). These MAPs with several possible origins preferentially derived from zinc finger proteins, which are known to share homologous regions (Supplementary Fig. S11). We therefore trained CAMAPs with datasets excluding entries encoding for MAPs that had >3 or >10 possible origins and compared their performance with that of CAMAPs trained without excluding these MAPs. Our results show that whatever the dataset used, CAMAP trained with original sequences always significantly outperformed CAMAP trained with shuffled sequences (Supplementary Fig. S12). Taken together, these results suggest that the codon-specific rules captured by CAMAP are independent of potential homologies in the hit dataset, as they do not appear to influence CAMAP performance.

We next validated our CAMAP trained on 9-mer MAPs derived from B-LCL using 5 datasets derived from different human and mouse cell types. All the validation datasets were described through proteogenomic analyses similarly to our B-LCL training datasets. However, all the validation datasets included MAPs of 8-11 mers, in contrast with the training dataset that contained only 9-mer MAPs. The validation datasets consisted of (i) our B-LCL dataset, this time including all peptide lengths [19,22], (ii) a dataset of human peripheral blood mononucleated cells or PBMCs [24], (iii) a dataset of B-lymphoblastoid cells expressing unique HLA alleles (B721.221 [11]), (iv) murine colon carcinoma cell line (CT26) and (v) a murine lymphoma cell line (EL4, [24,25]). For all datasets, we created hit and decoy datasets of original and shuffled sequences using the same approach described above but including MAPs of 8-11 amino acids. Notably, CAMAPs trained on human sequences encoding 9-mers MAPs from one human cell type (i.e. B-LCL) could also predict presentation of 8-11 mers MAPs in other human cell types (Fig. 4), as well as from mouse cell lines, albeit with lower performances (Fig. 4). Here again, CAMAPs trained on original sequences consistently outperformed CAMAPs trained on shuffled sequences (Fig. 4). These results show that the codon-specific rules derived by CAMAPs to predict MAP presentation are valid across different cell types, and can even be applied to another species, albeit with slightly lower performances. These results support a role for codons in the modulation of MAP presentation.

**Figure 4.**
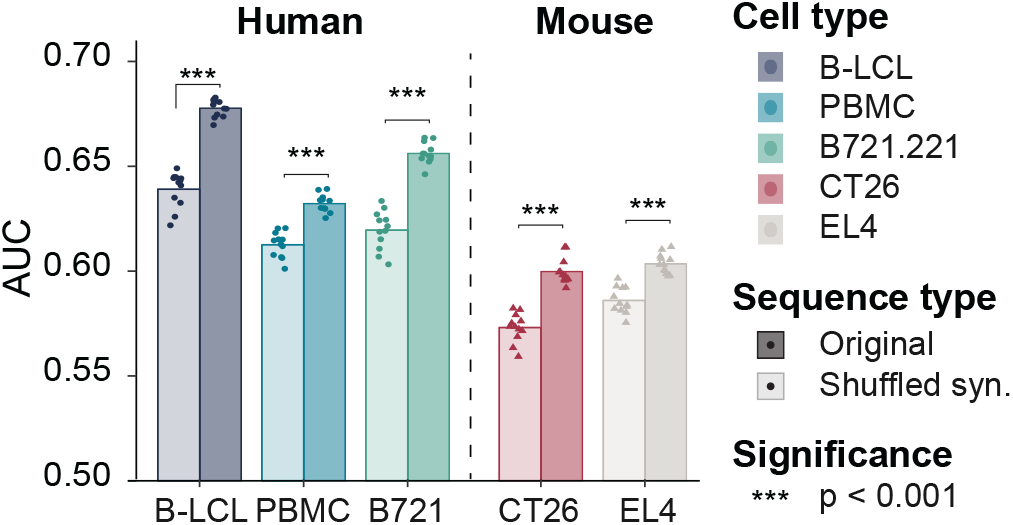
Validation of CAMAP predictions on 5 datasets derived from human and murine cell lines. CAMAP prediction score for different datasets derived from humans (i.e. B-LCL, PBMCs and B721.221) or mouse (i.e. CT26 and EL4) cells. Of note, all CAMAPs were trained on B-LCL-derived sequences encoding for 9-mer MAPs only with a context size of 162 nucleotides. Results are reported for 8 to 11-mer MAPs derived from the 5 datasets. In all panels, 12 CAMAPs trained with original or shuffled synonymous sequences were compared (significance assessed using Student T test).

The lower performances of CAMAP trained with shuffled sequences (representing amino acid distribution) suggests that amino acids in MAP-flanking sequences are less informative than codons regarding MAP presentation. We formally quantified this difference in information using the Kullback-Leibler (KL) divergence (see Materials and Methods for more details). Most codons (47/61, 77%) showed greater KL divergence in the original dataset than the shuffled dataset, indicating that codon distributions contained more information with regards to MAP presentation than amino acid distributions (Supplementary Fig. S13). These results suggest that codons in MAP-flanking regions play a role that is non-redundant with amino acids in MAP biogenesis.

We wondered whether some regions were more influential on MAP presentation than others. To address this question, we retrieved the model preferences for each codon at each position. The preferences correspond to the prediction score of our best model (trained with original codon sequences for a context size of 162 nucleotides) when a single codon at a single position is provided as input (all other positions being set at [0,0] coordinates in the embedding space). The model’s preferences are therefore a measure of each individual codon’s propensity to increase or decrease the model’s output probability as a function of its position relative to the MCCs. A value of 0.5 denotes a neutral preference, while negative and positive preferences correspond to values below and above 0.5, respectively. Preferences were obtained by feeding CAMAP sequences in which all codon values were masked, except for a single position that received a non-null codon label.

Interestingly, while codons closest to the MCCs were the most influential on CAMAP scores, some synonymous codons showed opposite effects, further demonstrating that codon usage does not recapitulate amino acid usage (Fig. 5A-B and Supplementary Fig. S14). The use of embeddings to encode codons has the advantage of arranging them into a semantic space, wherein codons with similar influences are positioned close to each other. Interestingly, most synonymous codons did not form clusters, with a notable exception being proline codons (Fig. 5C). This finding indicates that for some codons, their effect on CAMAP prediction score may be closer to that of a non-synonymous codon than to that of one of its synonyms.

**Figure 5.**
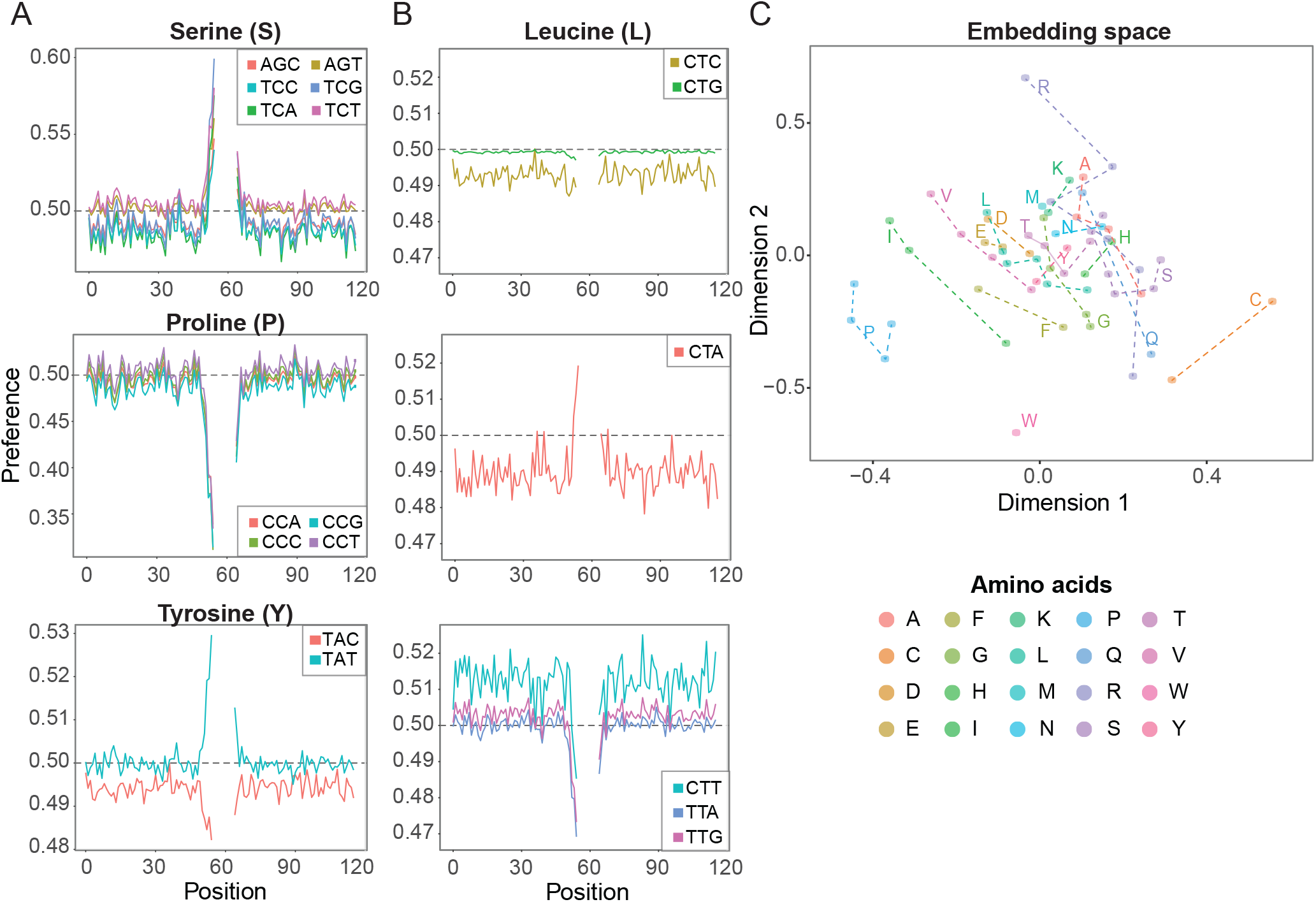
CAMAP interpretation of codon impact on MAP biogenesis. Preferences for a network trained on a context of 162 nucleotides (54 codons) for (A) serine, proline and tyrosine codons, and (B) leucine codons. (C) Learned codon embeddings. Some synonymous codons, such as those encoding for Isoleucine (I), Cysteine (C) or Arginine (R) are located far from one another, while others tend to cluster together (e.g. Proline [P] and Glutamic acid [E]).

### CAMAP increases MAP prediction accuracy

We next compared MAP prediction capacities of CAMAPs scores to that of MAP predicted ligand score (ranks as predicted by NetMHCpan4.0) and mRNA transcript expression levels. We used ligand scores as predicted by NetMHCpan4.0, which was shown to possess the best predictive capacities for naturally processed peptides compared to other predictive algorithms [26]. Because MAP binding to the MHC molecule is essential for its presentation at the cell surface, we elected to only compare hits and decoys encoding potential binders, i.e. with a minimal ligand score of 1% for at least one allele in the B-LCL dataset. Using a linear regression model, we compared the predictive capacity of each single parameter using Matthews correlation coefficient, which measures the quality of binary classifications [27]. Of note, only the predictions on the test set were used to evaluate the Matthew correlation coefficient in our different models.

Because only potential binders were analyzed here, the mRNA expression level had the highest predictive capacity, then followed by ligand scores (second) and CAMAP scores (third, Fig. 6A). As expected due to the multiplicative relationship between MAP ligand score and expression levels in predicting naturally processed MAPs [11], combining both variables greatly increased prediction performances (Fig. 6B). Importantly, adding CAMAP scores to the regression model further increased predictive performances (Fig. 6B). We next computed how many predicted peptides would need to be tested to capture 1, 5 or 10% of hits in the B-LCL dataset. Results presented in Table 1 show that using only NetMHCpan4.0 ligand scores (ranks) leads to a very high false positive rate (FPR) at 72.1% when targeting the top 1%. Adding the expression levels greatly increased prediction accuracy and decreased the FPR to 32.8% for the top 1% hits. When adding CAMAP scores as a third variable, the number of peptides needed to capture 1% of hits greatly decreased, resulting in a very low FPR at 1.1%. Similar trends were observed when targeting 5 or 10% of hits, although with higher FPR (see Table 1). Similarly, adding CAMAP scores to expression levels and ligand scores also ameliorated prediction accuracies for the two other human datasets introduced above (B721.221 and PBMCs, see Supplementary Table S2). These results show that combining CAMAP scores with the MAP’s ligand score (ranks) and its corresponding transcript expression level significantly improves prediction of MAP and facilitate identification of relevant epitopes through more accurate predictions.

**Figure 6.**
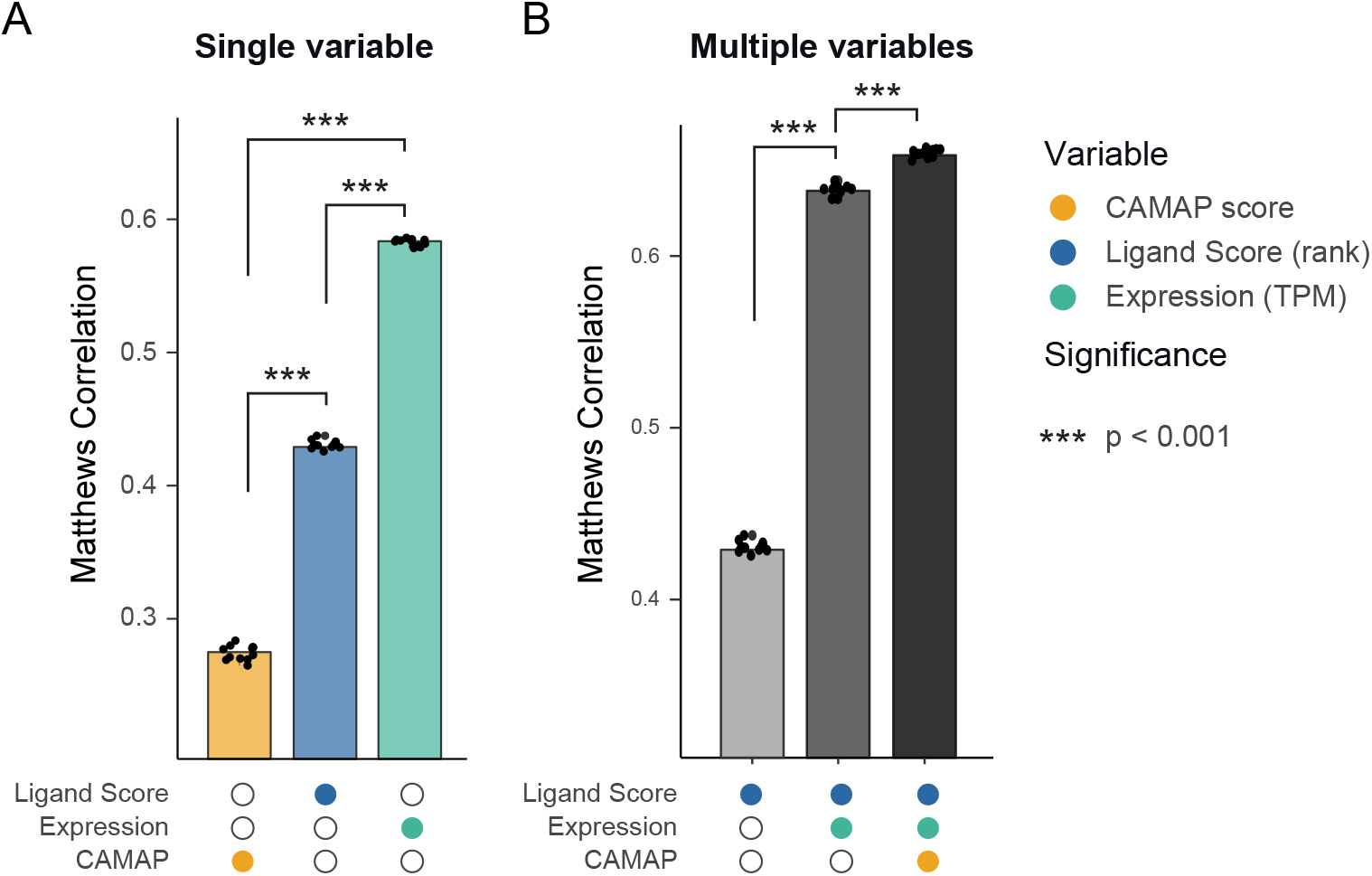
CAMAP prediction score contributes to the prediction of MAPs. (A) Matthews correlation coefficient for MAP prediction using a single variable. (B) Matthews correlation coefficient for MAP prediction using multivariable regression models. The B-LCL dataset (all MAP lengths) was filtered for MAP with a minimal ligand score (rank) of 1% (NetMHCpan4.0).

**Table 1.** Number of peptides needed to capture 1%, 5% or 10% of epitopes detected by mass spectrometry. The lower the number of peptides needed to capture the respective number of epitopes, the better the performance of the prediction model. This is also illustrated by the percentage of false identification (false positive rate, FPR) reported here. Peptides were rank-ordered according to regression scores, for a total of 490,297 unique peptides and 8,991 hits. Of note, only the maximal regression score was kept for peptides with multiple potential origins.

| Regression model | 1% hits (n=90) | 5% hits (n=450) | 10% hits (n=899) |
| --- | --- | --- | --- |

|  | n | FPR | n | FPR | n | FPR |
| --- | --- | --- | --- | --- | --- | --- |
| NetMHCpan4.0 | 322 ± 18 | 72.05% | 2183 ± 69 | 79.39% | 4927 ± 136 | 81.75% |
| NetMHCpan4.0<br>+ expression | 134 ± 6 | 32.84% | 601 ± 10 | 25.12% | 1211 ± 16 | 25.76% |
| NetMHCpan4.0<br>+ expression<br>+ CAMAP | **91 ± 2** | **1.11%** | **524 ± 13** | **14.12%** | **1170 ± 18** | **23.16%** |

### Codon usage can modulate MAP presentation

To evaluate whether changing the codon arrangement in a MAP-coding sequence might directly lead to modulation of MAP presentation, we generated three variants of the chicken ovalbumin (OVA) protein containing the model MAP SIINFEKL [28]. One construct encoded the wild type OVA (OVA-WT). For the other two constructs, we used CAMAP (trained on original human B-LCL sequences; Fig. 2) to generate two OVA variants *in silico*, both encoding for the same OVA protein but using different synonymous codons: one predicted to enhance SIINFEKL presentation (OVA-EP), the other predicted to reduce it (OVA-RP). Accordingly, the respective CAMAP scores for OVA-RP, OVA-WT and OVA-EP were: 0.03, 0.65, and 0.96 (Fig. 7A). All variants encoded the same amino acid sequence but used different synonymous codons. Notably, the sole difference between the three constructs were the 162 nucleotides flanking each side of the SIINFEKL-coding codons (i.e. the RNA sequences coding for OVA_202-256_ and OVA_265-319_, Supplementary Table S1 and Supplementary Figure S15).

**Figure 7.**
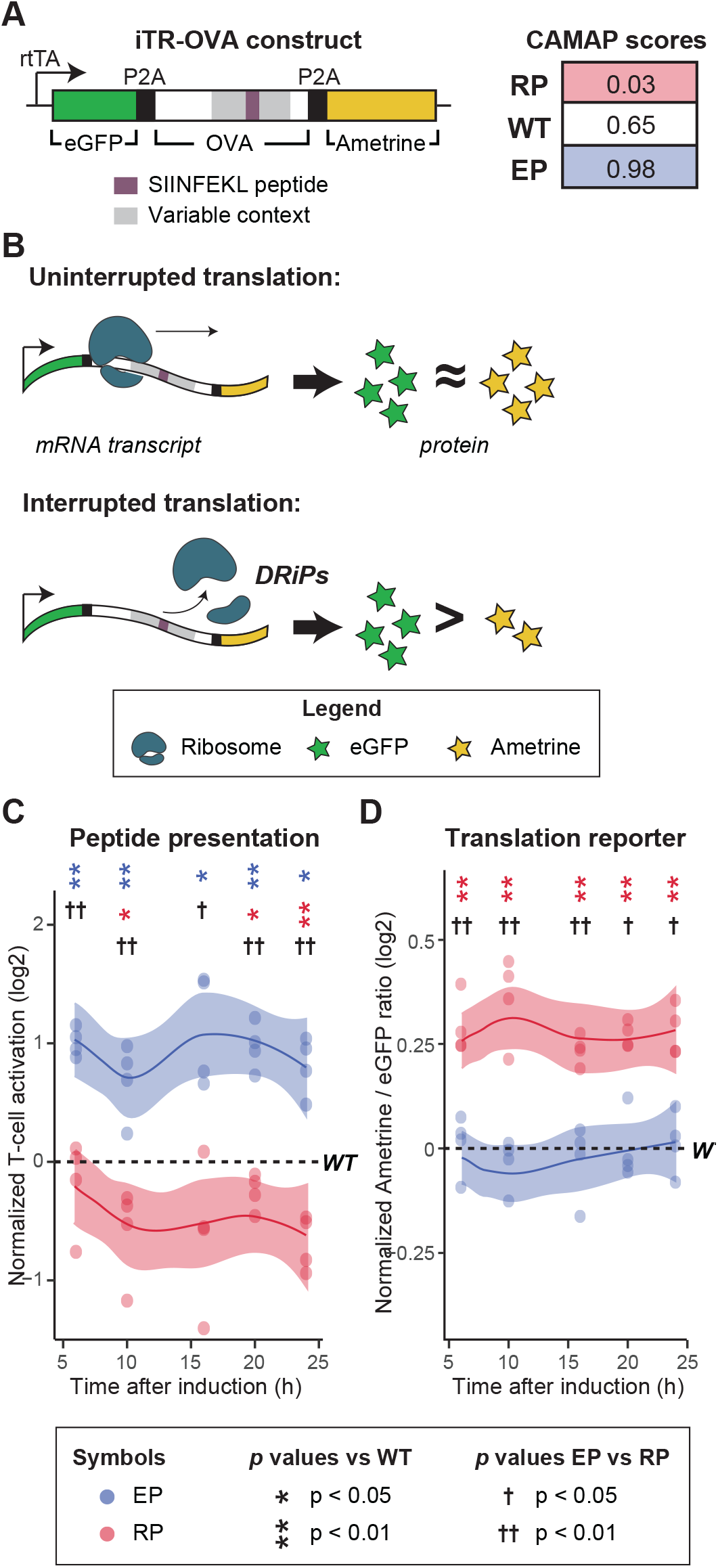
Codon usage in MAP-flanking mRNA sequences can influence antigen presentation and translation efficiency. (A) Design of the inducible Translation Reporter (iTR-OVA) constructs and CAMAP scores for OVA-WT, OVA-EP and OVA-RP sequences. (B) Schematic representation of possible translation events. When mRNA codon usage leads to efficient (uninterrupted) translation, similar amounts of eGFP and Ametrine proteins would be synthesized. When codon usage in the MAP-flanking regions enhances the frequency of translation interruption, a lower Ametrine/eGFP ratio would be observed. (C) Kinetics of SIINFEKL MAP presentation following induction of iTR-OVA constructs expression by doxycycline, measured in a T-cell activation assay. To remove the influence of differential expression levels on antigenic presentation and of varying proportion of transduced cells between samples, T-cell activation levels were normalized to the average Ametrine fluorescence intensity and to the proportion of eGFP+ cells (i.e. cells expressing the construct). (D) Translation efficiency as measured by Ametrine/eGFP ratio following iTR-OVA construct induction. For C and D, results are normalized over the WT sample from the same experiment (n=4). Statistical differences at each time point were determined using bilateral paired Student T tests. Significance for the comparison against WT are indicated with *, while comparison of EP vs RP is indicated with †. N.B.: Each replicate is shown with a dot, while the line and shaded area represent the average and 95% confidence interval, respectively.

Because codon usage affects translation efficiency, theoretically leading to DRiP formation through premature translation arrest [20,21], we expected the variable regions of our construct to affect both translation rates and SIINFEKL presentation in our variants. Therefore, each construct also coded for two other proteins, eGFP and Ametrine, placed upstream and downstream of the OVA coding sequence, respectively (Fig. 7A). While the Ametrine fluorescence intensity reflected the translation rate of the whole construct, the ratio of Ametrine/eGFP fluorescence intensity was informative regarding the translation efficiency of the whole construct. Indeed, efficient translation of the full-length construct should produce equivalent quantities of Ametrine and eGFP proteins, while inefficient/interrupted translation of the construct (i.e. leading to DRiP formation) should decrease the Ametrine/eGFP ratio (Fig. 7B). The three protein coding sequences were separated with P2A self-cleaving peptides [29], therefore allowing the co-synthesis of three separate proteins, controlled by the doxycycline-inducible Tet-On promoter. Importantly, the three proteins were tightly co-expressed because of the presence of only one start codon at the 5’ end of the GFP protein, as shown by the very high correlation between eGFP and Ametrine fluorescence for each construct (R>0.97, see Supplementary Figure S16). As we assumed that CAMAP scores reflected the probability of DRiP generation leading to increased MAP presentation, we expected the OVA-RP construct to show both reduced SIINFEKL presentation and enhanced translation efficiency compared to the OVA-EP and OVA-WT constructs. However, as both the OVA-EP and OVA-WT have CAMAP scores above the neutral threshold of 0.5 and closer to one another (0.98 and 0.65, respectively) compared to the OVA-RP construct (0.03), we expected OVA-EP and OVA-WT to behave more similarly.

We then used a SIINFEKL-H2-K^b^ specific T-cell activation assay [30] to measure SIINFEKL presentation at the cell surface following doxycycline induction. Results for the T-cell activation assay were normalized by both the Ametrine mean fluorescence intensity and the percentage of transduced (eGFP+) cells in each specific sample, so that any difference in T-cell activation observed between our constructs could only be ascribed to synonymous codon variants in the SIINFEKL-flanking OVA codons. Two main findings emerged from our analyses. First, in accordance with CAMAP predictions, variation in codon usage led to a 2.3-fold difference in SIINFEKL presentation between the OVA-EP and OVA-RP variants, with OVA-WT in between (Fig. 7C). Second, translation efficiency (Ametrine/eGFP ratio) was higher with OVA-RP than with OVA-EP or OVA-WT, while OVA-EP showed similar translation efficiency compared to ONA-WT (Fig. 7D). Hence, synonymous codon variations led to slightly divergent outcomes in OVA-EP and OVA-RP: they modulated the levels of SIINFEKL presentation in both constructs, but enhanced translation efficiency could only be detected for OVA-RP. These data show that codon arrangement can modulate MAP presentation strength without any changes in the amino acid sequence and support a role for translation efficiency and DRiP formation in the modulation of MAP presentation.

## Discussion

Our analyses of large datasets using artificial neural networks and other bioinformatics approaches provide compelling evidence that codon usage regulates MAP biogenesis via both short- and long-range effects. While most MAP predictive approaches focus on MAP sequences *per se*, CAMAP’s novelty is that it only receives the MAP-flanking mRNA sequences as input, and no information on the MAP itself, thereby providing completely independent information for MAP prediction. The better prediction accuracy of CAMAPs trained with original codons rather than with shuffled synonyms supports the role of codon usage in modulating MAP biogenesis (Fig. 2). In addition, we demonstrated that the codon-specific signal that is captured by CAMAP was independent of transcript expression levels and MAP ligand scores, thereby providing complementary and non-overlapping information regarding MAP presentation. Additionally, while CAMAP preferences were more influential for codons located close to the MCCs (Fig. 5), the better performance of CAMAP trained with longer context size pointed toward a long-range impact of codon usage on MAP presentation.

The functional link between codon arrangement and MAP biogenesis was illustrated by our *in vitro* analyses of SIINFEKL biogenesis, in which we were able to modulate SIINFEKL presentation solely by substituting synonymous codons in mRNA regions flanking SIINFEKL codons, without changing the protein sequence. While the experimental data derives from a single model thus limiting the interpretability of our results, this points nonetheless to an interesting mechanism that could be exploited to enhance antigenic presentation in peptide-bas4ed immunotherapy (i.e. dendritic cells modified to express a specific MAP).

Further analyses will be needed to assess the full extent of codon arrangement’s impact on both classic MAPs (i.e. derived from canonical reading frames of coding sequences) and cryptic MAPs (i.e. derived from non-canonical reading frames and non-coding sequences) [31,32], as well as the potential contribution of codons in non-coding regions (e.g. 5’- or 3’-UTRs) on the regulation of MAP presentation. However, our results show that the integration of CAMAP scores to the two best predictive factors for naturally processed MAPs led to a significant increase in prediction accuracy. Indeed, our regression model combining only transcript expression levels to MAP ligand scores (ranks as predicted by NetMHCpan4.0), showed that a total of 134 peptides would need to be tested in order to capture 1% of all presented MAPs (hits), leading to a false positive rate of 32.8%. In contrast, the addition of CAMAP to this model decreased the false positive rate to only 1.1%, leading to 90 correct identifications out of 91 MAPs tested. Although predictions were not as accurate for the two other human datasets, adding CAMAP scores always resulted in improved prediction accuracy. Our results therefore support the combined use of ligand scores, transcript expression levels and CAMAP scores in MAP predictive algorithms. These results have important practical implications for cancer immunotherapy and peptide-based vaccines, where discovery of suitable target antigens remains a formidable challenge to this day [33,34].

## Materials and methods

### Dataset generation

We analyzed a previously published dataset consisting of MAPs presented on B lymphocytes by a total of 33 MHC-I alleles from 18 subjects [19,22]. Since this dataset was assembled using older versions of MHC-I binding prediction algorithms (i.e. using a combination of NetMHC3.4 for common alleles and NetMHCcons1.1 for rare alleles), we verified that the majority of MAPs in this dataset would also be predicted as binders using more recent algorithms (i.e. a rank ≤ 2.0% using NetMHC4.0 or NetMHCpan4.0). We found an overlap of >92% between these methods (see Supplementary Fig. S17), thereby validating this dataset for further analysis. In addition, we reasoned that a transcript should be considered as a genuine positive or negative regarding MAP biogenesis only if it was expressed in the cells. We therefore excluded from the dataset all transcripts with very low expression (<1^st^ percentile in terms of FPKM).

To facilitate data analysis and interpretation, we only included transcripts coding for MAPs with a length of 9 amino acids, for a total of 19,656 9-mer MAPs (which represents 78% of MAPs in this dataset). We then used pyGeno [23] to extract the mRNA sequences of transcripts coding for these 9-mer MAPs, which constituted our source-transcripts (Fig. 1A). We next created a negative (non-source) dataset from transcripts that generated no MAPs. Importantly, transcripts that encoded for MAPs of any length (i.e. 8 to 11-mer) were excluded from the negative dataset. We then randomly selected 98,290 non-MAP 9-mers from this negative dataset, and extracted their coding sequences using pyGeno. Of note, both positive and negative datasets were derived from the canonical reading frame of non-redundant transcripts.

We analyzed only the MAP context and excluded the MCCs *per se* from our positive (hits) and negative (decoys) sequences (Fig. 1A). We limited our analyses of flanking sequences to 162 nucleotides (54 codons) on each side of MCCs, because longer lengths would entail the exclusion of >25% of transcripts (Supplementary Fig. S18).

### Creation of the shuffled synonymous codon dataset

To create the shuffled synonymous codon dataset, each sequence was re-encoded by replacing each codon with itself or with a random synonym according to the human transcriptome usage frequencies. These frequencies were calculated using the annotations provided by *Ensembl* for the human reference genome GRCh37.75. Thus, all codon-specific features differing between the positive and negative datasets was removed from the shuffled datasets. Because codons were replaced by their synonymous codons, the shuffled sequences directly reflected amino acid usage in the positive and negative datasets.

### CAMAP architecture, sequence encoding and training

The first (input) layer received either MCCs flanking regions from the hit dataset or sequences of the same length contained in the decoy dataset (Fig. 1A). The second layer (Supplementary Fig. 4A) was a codon embedding layer similar to that introduced for a neural language model [35]. Embedding is a technique used in natural language processing to encode discrete words, and has been shown to greatly improve performances [36]. With this technique, the user defines a fixed number of dimensions in which words should be encoded. When the training starts, each word receives a random vector-valued position (its embedding coordinates) in that space. The network then iteratively adjusts the words’ embedding vectors during the training phase and arranges them in a way that optimizes the classification task. Notably, embeddings have been shown to represent semantic spaces in which words of similar meanings are arranged close to each other [36]. In the present work, we treated codons as words: each codon received a set of random 2D coordinates that were subsequently optimized during training. The third (output) layer delivered the probability that the input sequence was a MCCs flanking region (rather than a sequence from the negative dataset).

CAMAPs were trained on sequences resulting from the concatenation of pre- and post-MCCs regions. Before presenting sequences to our CAMAPs, we associated each codon to a unique number ranging from 1 to 64 (we reserved 0 to indicate a null value) and used this encoding to transform every sequence into a vector of integers representing codons. Neural networks were built using the Python package Mariana [37] [https://www.github.com/tariqdaouda/Mariana]. The *Embedding* layer of Mariana was used to associate each label superior to 0 to a set of 2D trainable parameters; the 0 label represents a *null* (masking) embedding fixed at coordinates (0,0). As an output layer, we used a *Softmax* layer with two outputs (positive / negative). Because negative sequences are more numerous than positive ones, we used an oversampling strategy during training. At each epoch, CAMAPs were randomly presented with the same number of positive and negative sequences. All CAMAPs in this work share the same architecture (Supplementary Fig. 4A), number of parameters and hyper-parameter values: learning rate: 0.001; mini-batch size: 64; embedding dimensions: 2; linear output without offset on the embedding layer; *Softmax* non-linearity without offset on the output layer.

For each condition (e.g. context size), the positive and negative datasets were randomly divided into three non-redundant subsets: (i) the training subsets containing 60% of the positive and negative transcripts, (ii) the validation and (iii) the test subsets each containing 20% of the positive and negative transcripts. Transcripts were assigned through a sequence redundancy removal algorithm, thereby ensuring that no transcript was assigned to multiple subsets. We used an early stopping strategy on validation sets to prevent over-fitting and reported average performances computed on test sets. We trained 12 CAMAPs for each combination of conditions, each one using a different random split of train/validation/test sets. To mask sequences either before or after the MCCs, we masked either half with *null* value.

### Kullback-Leibler divergence

The Kullback-Leibler (KL) divergence computes how well a given distribution is approximated by another distribution. Its value can be either positive or 0, a null value indicating that the two distributions are identical (see Materials and Methods for more details). Accordingly, a higher KL divergence for codon distributions vs. amino acid distributions would indicate that codon variations are not entirely accounted for by amino acid variations. KL divergence is not a metric, as it is neither symmetric nor does it satisfy the triangle inequality. It is nevertheless an accurate and most common way of comparing two probability distributions.

We defined the probability of having codon *c* at position *i* as a function of the number of occurrences of *c* at position *i*, divided by the total number of occurrences of that same codon:

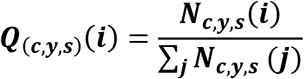

Here Q is a probability, N is a number of occurrences, c is a codon, y is a class (positive or negative), s indicates if codons have been randomized (true or false), i is a position in sequence. For the remainder of the text we will use the following abbreviations:

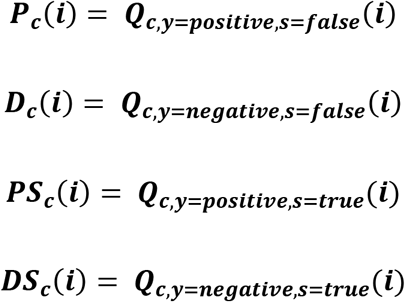

We then used the KL divergence to compute how well *P*_*c*_ distributions approximate *D*_*c*_ distributions and *PS*_*c*_ distributions approximate *DS*_*c*_ distributions.

The KL divergence was defined as:

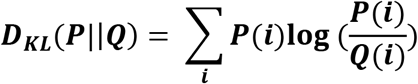

We performed this calculation for both the original and the shuffled dataset, which we then compared together. If codons and amino acid distributions were equivalent, KL divergence between hits and decoys would be the same for both original and shuffled sequences, and codons would cluster along the diagonal.

### Predicting MAP presentation with linear regressions

The prediction capacity of CAMAP, NetMHCpan-4.0 ligand score and transcription expression (TPM) was tested in different combinations of those parameters (Ligand Score + Expression, Ligand score + Expression + CAMAP score) using the *LogisticRegressionCV* function from the python package *sklearn* (*sklearn.linear_mode*l, v0.22.1). In each case, the dataset containing hits and decoy sequences was split into train and test datasets with a ratio of 0.7 to 0.3, respectively. Values for CAMAP score, Ligand Score and TPM were each scaled to a range of 0-1 in the train set using MinMaxScaler from *sklearn.preprocessing* and the same scaling model was applied to the test set afterwards. Regression analysis was performed using *LogisticRegressionCV* with a 10x cross-validation using the *lbfgs* solver with 1000 iterations. MCC scores were calculated using *matthews_corrcoef* from *sklearn.metrics*. When a peptide had multiple sources (multiple transcripts or genes), only the maximum value from its regression scores was kept.

### In vitro assay – inducible translation reporter (iTR)-OVA construct design

An inducible translation reporter was generated by flanking the truncated chicken ovalbumin (OVA) cDNA (amino acids 144-386) with EGFP-P2A (in 5’) and P2A-Ametrine (in 3’) cDNA sequences. MCCs flanking contexts for the EP and RP construct were synthesized as gBlocks (purchased from Integrated DNA Technologies). The fragments were amplified by PCR and joined by Gibson assembly under a doxycycline-inducible Tet-ON promoter in a pCW backbone. Synthetic variants of the OVA coding sequence were generated in silico by varying synonymous codon usage in the MAP context regions (i.e. 162 nucleotides pre- and post-MCCs). Importantly, the amino acid sequence was preserved between the different variants; only nucleotide sequences in the MAP context (162 nucleotides on either side) differed. The sequences with the highest (EP) and the lowest (RP) prediction scores were selected for further in vitro validation and swapped into the iTR-OVA plasmid by Gibson assembly [38]. OVA-EP and OVA-RP sequences can be found in Supplementary Table 1.

Important features of our inducible translation reporter construct and T cell activation assay were: (i) No changes in amino acid sequence between the three variants: only co-translational events can differ between the three variants, post-translational events being equivalent for the three constructs; (ii) Only one start codon, at the beginning of the eGFP coding sequence: this is important for the translation reporter aspect of our construct (i.e. Ametrine/eGFP ratio), to ensure that translation can only start at the 5’-end of the whole construct, and not at the beginning of the OVA or Ametrine coding sequences; (iii) Separation of the three proteins using P2A peptide: allows the inducible synthesis of three separate proteins in a highly correlated manner; also, the degradation of one protein will be independent from the others. As we hypothesized that codon usage might lead to DRiP formation, we did not want the degradation of OVA-derived polypeptide to induce degradation of attached eGFP or Ametrine, which would affect our translation reporter assay (Ametrine/eGFP ratio); (iv) Because transcript expression level impacts MAP presentation, we normalized T-cell activation results by both the number of transduced cells present in the samples (% of eGFP+ cells) and the Ametrine mean fluorescence intensity of eGFP+ cells (representing whole construct expression level). Because of these four features, any difference between the three constructs could be ascribed solely to synonymous codon variants in the SIINFEKL-flanking OVA codons.

### Stable cell line generation

Wildtype and transduced Raw-K^b^ cells [39] were cultured in DMEM supplemented with 10% Fetal Bovine Serum (FBS), penicillin (100 units/ml), and streptomycin (100mg/ml). B3Z cells [40] were maintained in RPMI medium supplemented with 5% FBS, penicillin (100 units/ml), and streptomycin (100mg/ml).

Lentiviral particles were produced from HEK293T cells by co-transfection of iTR-OVA WT, EP or RP along with pMD2-VSVG, pMDLg/pRRE and pRSV-REV plasmids. Viral supernatants were used for Raw-K^b^ transduction. Raw-K^b^ OVA-WT, Raw-K^b^ OVA-EP were sorted on Ametrine and GFP double positive population after 24h of doxycycline treatment (1 mg/ml).

### T-cell activation assay

Raw-K^b^ OVA-EP, OVA-RP and OVA-WT cells were plated at a density of 250,000 cells/well in 24 well-plates 24h prior to doxycycline treatment (1 mg/ml). After the corresponding treatment duration, cells were harvested and fixed using PFA 1% for 10 minutes at room temperature and washed using DMEM 10% FBS. Raw-K^b^ were then co-cultured (37°C, 5% CO_2_) in triplicates with the CD8 T cell hybridoma cell line B3Z cells at a 3:2 ratio for 16h (7.5 × 10^5^ B3Z and 5 × 10^5^ Raw-K^b^) in 96 well-plates. Cells were lysed for 20 minutes at room temperature using 50 μl/well of lysis solution (25mM Tris-Base, 0.2 mM CDTA, 10% glycerol, 0.5% Triton X-100, 0.3mM DTT; pH 7.8). 170 μl/well CPRG buffer was added (0.15mM chlorophenol red-β-d-galactopyranoside (Roche), 50mM Na_2_HPO_4_•7H_2_0, 35mM NaH_2_PO_4_•H_2_0, 9mM KCl, 0.9mM MgSO_4_•7H_2_O). β-galactosidase activity was measured at 575 nm using SpectraMax^®^ 190 Microplate Reader (Molecular Devices). In parallel, cells were analyzed by flow cytometry using a BD FACS CantoII for eGFP and Ametrine fluorescence.

## Abbreviations

MHC-I: major histocompatibility complex class-I
MAP: MHC-I associated peptides
CAMAP: Codon arrangement MAP predictor
DRiP: defective ribosomal product
ANN: artificial neural network
MCC: MAP-coding codons
B-LCL: B-lymphoblastoid cell line
KL: Kullback-Leibler
BS: binding score
OVA: ovalbumin protein
WT: wildtype
EP: enhanced presentation
RP: reduced presentation

## Data Availability

The datasets analyzed for this study can be found:

- <u>Human B-LCL</u>: RNA-Seq data can be accessed on the NCBI Bioproject database (http://www.ncbi.nlm.nih.gov/bioproject/; accession PRJNA286122).
- <u>Human PBMC</u>: RNA-sequencing data for human PBMC were extracted from healthy donors in Zucca et al (2019) [41] and can be accessed under the GEO accession number GSE106443 and GSE115259, while MAPs were extracted from Murphy et al (2017) [24].
- <u>Human B721.221</u>: The B721.221 dataset was retrieved from Abelin et al (2017) [11]; RNA sequencing data can be accessed under the GEO accession number GSE93315.
- <u>Murine CT26</u>: RNA-Seq data can be accessed under the GEO accession number GSE111092. Mass spectrometry data can be found on the ProteomeXchange Consortium via the PRIDE partner repository (human B-LCL: PXD004023 and murine CT26: PXD009065 and 10.6019/PXD009065).
- <u>Murine EL4</u>: MAP dataset was extracted from Murphy et al (2017) [24] and EL4 RNA sequencing dataset was extracted from Sidoli et al (2019) [42] and can be accessed under the GEO accession number GSE125384.

All figures were generated using R’s package “ggplot2”. Source code for pyGeno (https://github.com/tariqdaouda/pyGeno, doi: 10.12688/f1000research.8251.2) and Mariana (https://github.com/tariqdaouda/Mariana, doi: [to be provided after acceptance]) are freely available online.

## Acknowledgements

This work was supported by grants from the Canadian Cancer Society (number 705604 and 705714), the Oncopole and the Leukemia & Lymphoma Society of Canada. Perreault’ lab is supported in part by The Katelyn Bedard Bone Marrow Association. MDL was supported by a studentship from the Canadian Institute of Health Research. Y.Benslimane was supported by a fellowship from the Cole Foundation and a Canadian Institutes of Health Research operating grant (to LH: #13784). EG lab is supported by the Canadian Institute for Health Research operating grant (MOP-133726). The B3Z CD8+ T cell hybridoma cell line was a kind gift from Nilabh Shastri. RAW-Kb cells were kindly provided by Michel Desjardins.

## Author contributions

TD designed all computational experiments. TD and AF performed computational experiments. TD wrote pyGeno and Mariana, contributed to design of the iTR-OVA construct, co-wrote the first draft of the paper. MDL contributed to data analysis, to design and synthesis of the iTR-OVA construct, performed flow cytometry analysis, with input of EG, co-wrote the first draft of the paper. AF contributed to data analysis, study design and performed computational experiments (validation on 5 datasets and regressions). Y.Benslimane contributed to design and synthesis of the iTR-OVA construct, with input from LH and EG. RP produced viruses for transduction of the iTR-OVA construct, transduced RAW cells, optimized and performed T-cell activation assay using mild fixation, with input from EG, and reviewed the manuscript. MC performed peptide affinity predictions. MB contributed to the optimization of culture conditions for the iTR-OVA assay. PT reviewed the manuscript. Y.Bengio reviewed and contributed to the manuscript. SL and CP contributed to study design, reviewed and contributed to the manuscript. All co-authors reviewed the manuscript.

The authors declare no competing interests.

## Supporting information captions

### Supplementary Figures

**Supplementary Figure S1. Codon distribution in the shuffled datasets more closely resembles that of amino acids, compared to the original datasets.** (A) Pearson correlation (R^2^) factors and (b) Kullback-Leibler (KL) divergence between positional distribution of codons and their corresponding amino acid in the shuffled (y axis) VS original (x axis) datasets. For all codons, the shuffled dataset showed greater correlations (A) and smaller KL divergence to their respective amino acid distributions than the original datasets (p < 1 x 10^−8^, assessed using unilateral paired Student T test).

**Supplementary Figure S2. Distribution of Pearson’s correlation factors calculated between codons and amino acids positional distributions in the original (green) and shuffled (coral) datasets.** 92% of codons in the shuffled dataset reflecting the amino acids distribution with a R2 > 0.95, compared to only 69% in the original dataset (p < 5×10^−5^).

**Supplementary Figure S3. Distribution of amino acid and codon usage per position in the original VS shuffled datasets.** (A) Alanine – A. (B) Cysteine – C. (C) Aspartic acid – D. (D) Glutamic acid – E. (E) Phenylalanine – F. (F) Glycine – G. (G) Histidine – H. (H) Isoleucine – I. (I) Lysine – K. (J) Leucine – L. (K) Asparagine – N. (L) Proline – P. (M) Glutamine – Q. (N) Arginine – R. (O) Serine – S. (P) Threonine – T. (Q) Valine – V. (R) Tyrosine – Y.

**Supplementary Figure S4.** CAMAP architecture and detailed predictions. (A) Architecture of the ANN used in this work. (B) Results for the AUC on all train, validation and test subsets. Grey areas represent the 95% confidence intervals. (C) Distributions of output probabilities of CAMAPs used to calculate correlations in Supplementary Figure S5.

**Supplementary Figure S5**. Correlation between CAMAP prediction score trained only with pre-MCC or post-MCC sequences. For each sequence in the test set we calculated the average prediction score given by CAMAPs in each condition, and calculated the Pearson correlation using the R software. Densities were calculated on all points and drawn using ggplot2. Only a random subset of the points is represented in the figures to limit their size.

**Supplementary Figure S6.** Absence of correlation between CAMAP prediction score and transcript expression levels in 4 individual B-LCL samples (each derived from a different subject).

**Supplementary Figure S7. Training of CAMAP on dataset selected to reflect positive dataset’s distribution in expression levels.** (A) Distribution of transcript expression levels for normal datasets (related to Figure 2) and the dataset used here to retrain CAMAP. As shown in this figure, the decoy dataset was selected to mirror the distribution of transcript expression level in the hit dataset. (B) CAMAP performance (measured by the AUC) when trained using the decoy dataset that mirrors the transcript expression levels of the hit dataset. Significance was assessed using bilateral paired Student T test (*p* = 5.36 x 10^−7^).

**Supplementary Figure S8. Absence of correlation between CAMAP prediction score and binding affinities for individual alleles for decoys (A) and hits (B).**

**Supplementary Figure S9. Training of CAMAP on dataset selected to reflect positive dataset’s distribution in binding affinities.** (A) Distribution of binding affinities for normal datasets (related to Figure 2) and the corrected dataset used to retrain CAMAP. As shown in this figure, the decoy dataset was selected to mirror the distribution of binding affinities in the hit dataset. (B) CAMAP performance (measured by the AUC) when trained using the decoy dataset that mirrors the binding affinities of the hit dataset. Significance was assessed using bilateral paired Student T test (*p* = 1.21 x 10^−9^).

**Supplementary Figure S10. Evaluation of homology in hit dataset and its impact of CAMAP performance.** (A) Proportion of unique MAPs that can be ascribed to a single origin, 2-3, or >10 possible origins. (B) Proportion of entries in the hit dataset that encode for MAPs with a single origin, 2-3, 4-10 or >10 possible origins

**Supplementary Figure S11. Gene families overrepresented in hits with >3 possible origins.**

**Supplementary Figure S12. CAMAP performance (AUC) when trained using either all hits (left), hits with 10 possible origins or less (center) or hits with 3 possible origins or less (right)**.

**Supplementary Figure S13**. Kullback-Leibler divergence between hit and decoy datasets in original codon (y-axis) or shuffled synonymous codon sequences (x-axis). Shuffled sequences represent amino acid usage, as codon-specific information are removed with synonymous codon shuffling.

**Supplementary Figure S14. Preferences per position for all codons for CAMAP trained with original sequences.** See Materials and Methods for more details.

**Supplementary Figure S15.** OVA-construct alignment, showing point mutations (red lines) in the mRNA sequences flanking the SIINFEKL MCC. (A) Comparison of the OVA-EP nucleotide sequence to the wildtype OVA sequence. The OVA-EP and OVA-WT sequences have 93.3% nucleotide identity for a total of 78 modified nucleotides. (B) Comparison of the OVA-RP nucleotide sequence to the wildtype OVA sequence. The OVA-EP and OVA-WT sequences have 92.6% nucleotide identity, for a total of 86 modified nucleotides. Mutations, shown in red, are located only in the 162 nucleotide regions flanking the SIINFEKL coding codons. Of note, the SIINFEKL coding codons (nucleotides 772-799) were not modified between the 3 constructs.

**Supplementary Figure S16.** Correlations between eGFP and Ametrine fluorescence intensity at the single cell level. Single cell eGFP and Ametrine fluorescence intensities measured at 10 hours post-induction are shown for the OVA-WT (A), OVA-EP (B) and OVA-RP (C) constructs. N.B.: only transduced cells are shown (eGFP+ cells).

**Supplementary Figure S17.** Validation of MHC-I associated peptides (MAP) dataset from Pearson H. *et al*. (2016) using the new versions of MAP binding affinity prediction algorithm NetMHC4.0 (A) and NetMHCpan4.0 (B).

**Supplementary Figure S18.** Percentage of transcript ineligibility as a function of context size. Transcript length corresponds to *C x 2 + 27*, where *C* is the context size in nucleotides and 27 the length of the MCCs. Related to Figure 1A.

### Supplementary Tables

**Supplementary Table S1. Nucleotide sequences of the EP and RP constructs.** SIINFEKL MCCs are shown in bold, while the variant regions (pre- and post-MCCs flanking sequences, context size of 162-nucleotides) are in blue and italics. Related to Fig. 7.

**Supplementary Table S2. Number of peptides needed to capture 1%, 5 and 10% of epitopes detected by mass spectrometry in B721.221 and PBMC cell lines.** The lower the number of peptides needed to capture the respective number of epitopes, the better the performance of the prediction model. This is also illustrated by the percentage of false identification (false positive rate, FPR) reported here. Peptides were rank-ordered according to regression scores. Of note, only the maximal regression score was kept for peptides with multiple potential origins.

