## Supplemental information for "CAMAP: Artificial neural networks unveil the role of codon arrangement in modulating MHC-I peptides presentation"

### Table of Contents

Supplementary Figures 3

Supplementary Tables 8

### Supplementary Figures

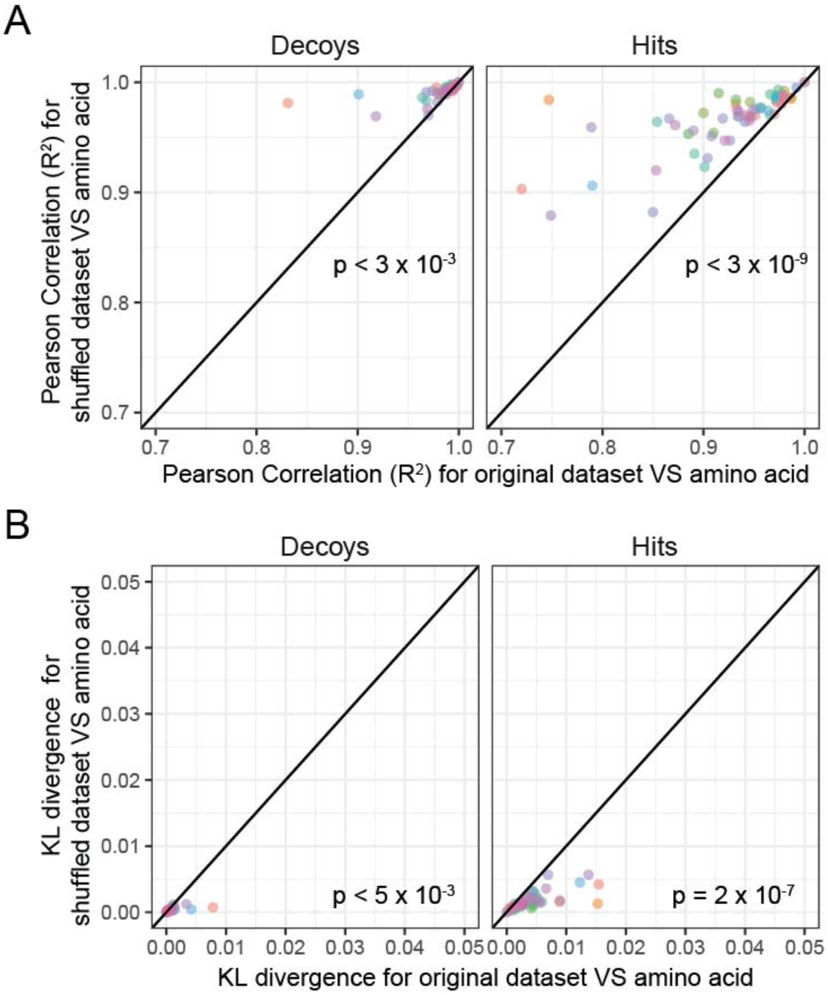


**Supplementary Figure S1. Codon distribution in the shuffled datasets more closely resembles that of amino acids, compared to the original datasets.** (A) Pearson correlation ($R^2$) factors and (b) Kullback-Leibler (KL) divergence between positional distribution of codons and their corresponding amino acid in the shuffled (y axis) VS original (x axis) datasets. For all codons, the shuffled dataset showed greater correlations (A) and smaller KL divergence to their respective amino acid distributions than the original datasets ($p < 1 \times 10^{-8}$, assessed using unilateral paired Student T test).

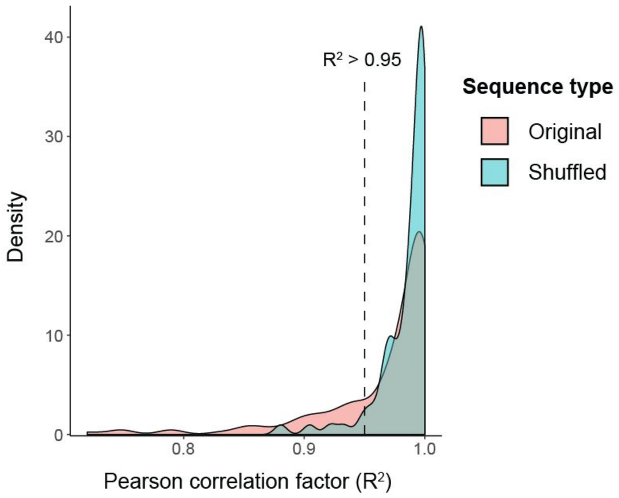


**Supplementary Figure S2. Distribution of Pearson's correlation factors calculated between codons and amino acids positional distributions in the original (green) and shuffled (coral) datasets.** 92% of codons in the shuffled dataset reflecting the amino acids distribution with a R2 > 0.95, compared to only 69% in the original dataset ($p < 5x10^{-5}$).

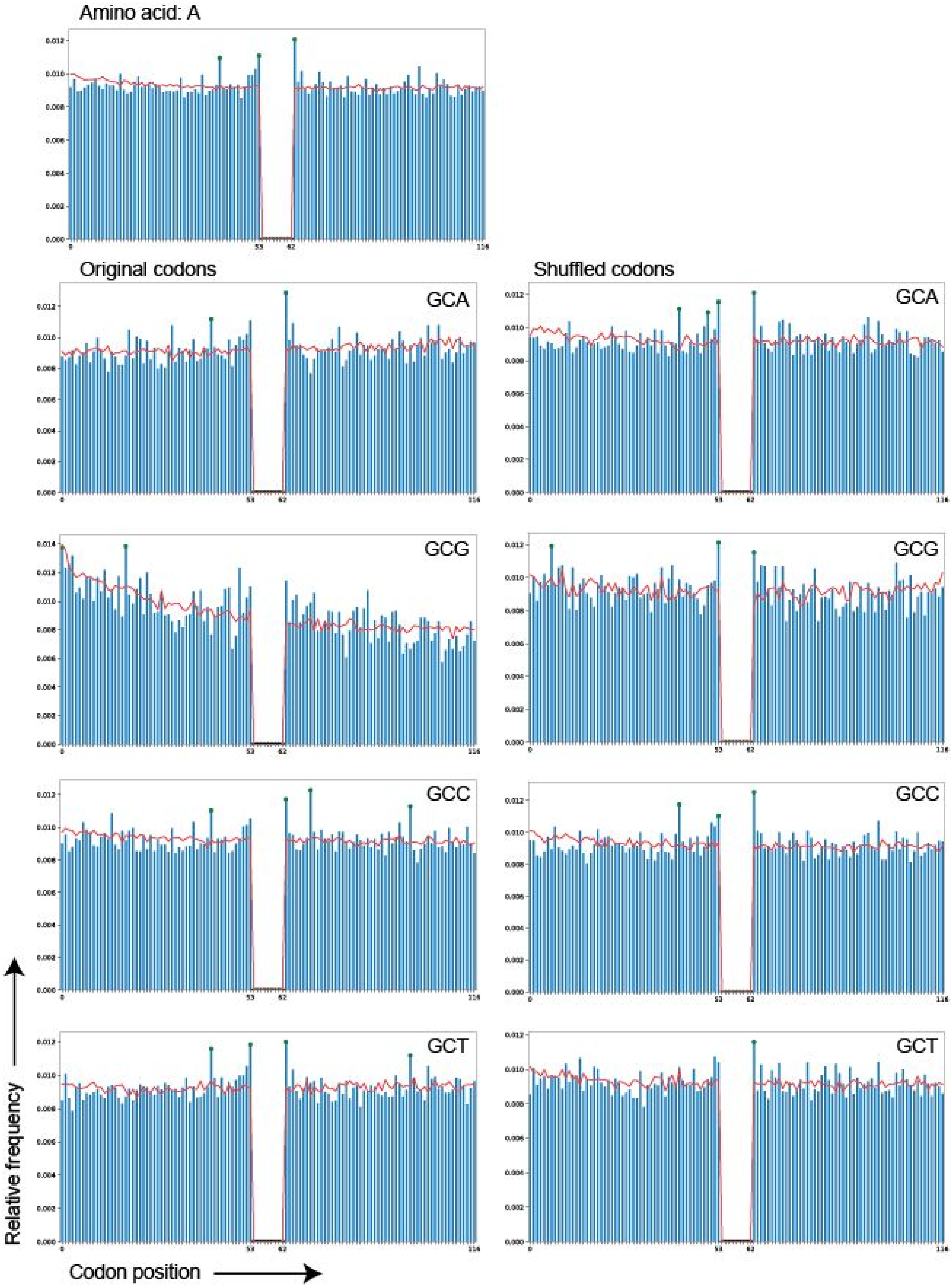


**Supplementary Figure S3. Distribution of amino acid and codon usage per position in the original VS shuffled datasets.** (A) Alanine (A).

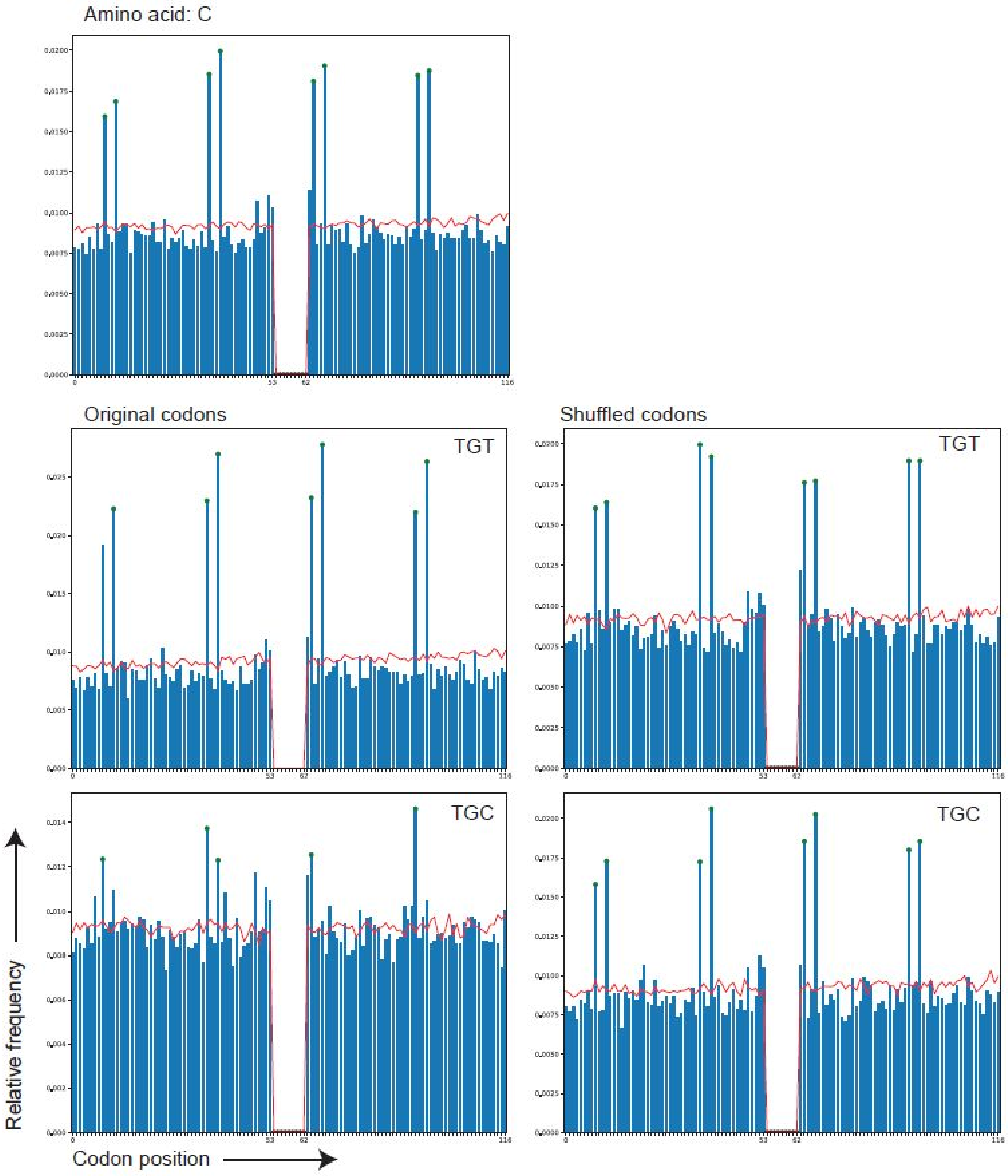


**Supplementary Figure S3. Continued.** (B) Cysteine (C).

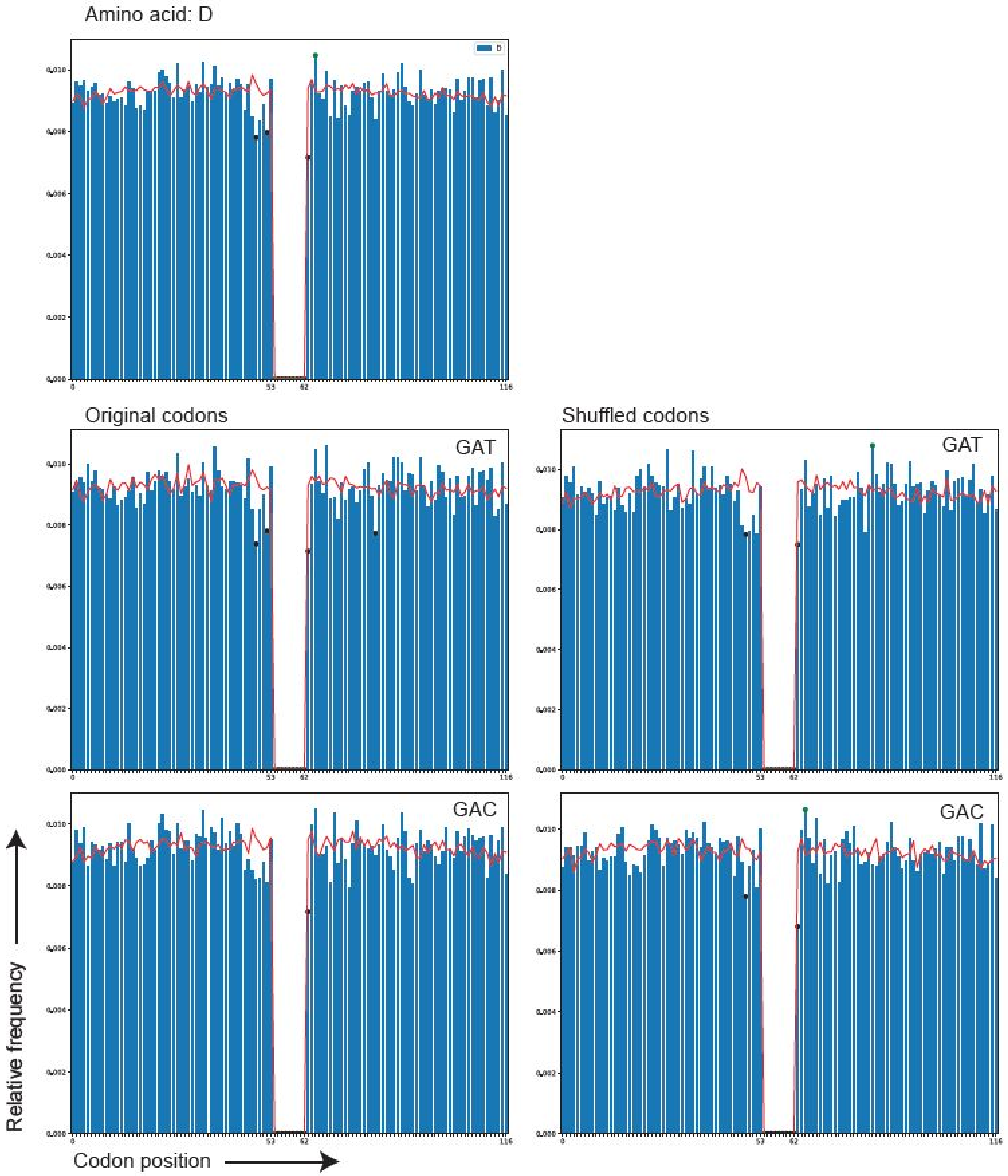


**Supplementary Figure S3. Continued.** (C) Aspartic acid (D).

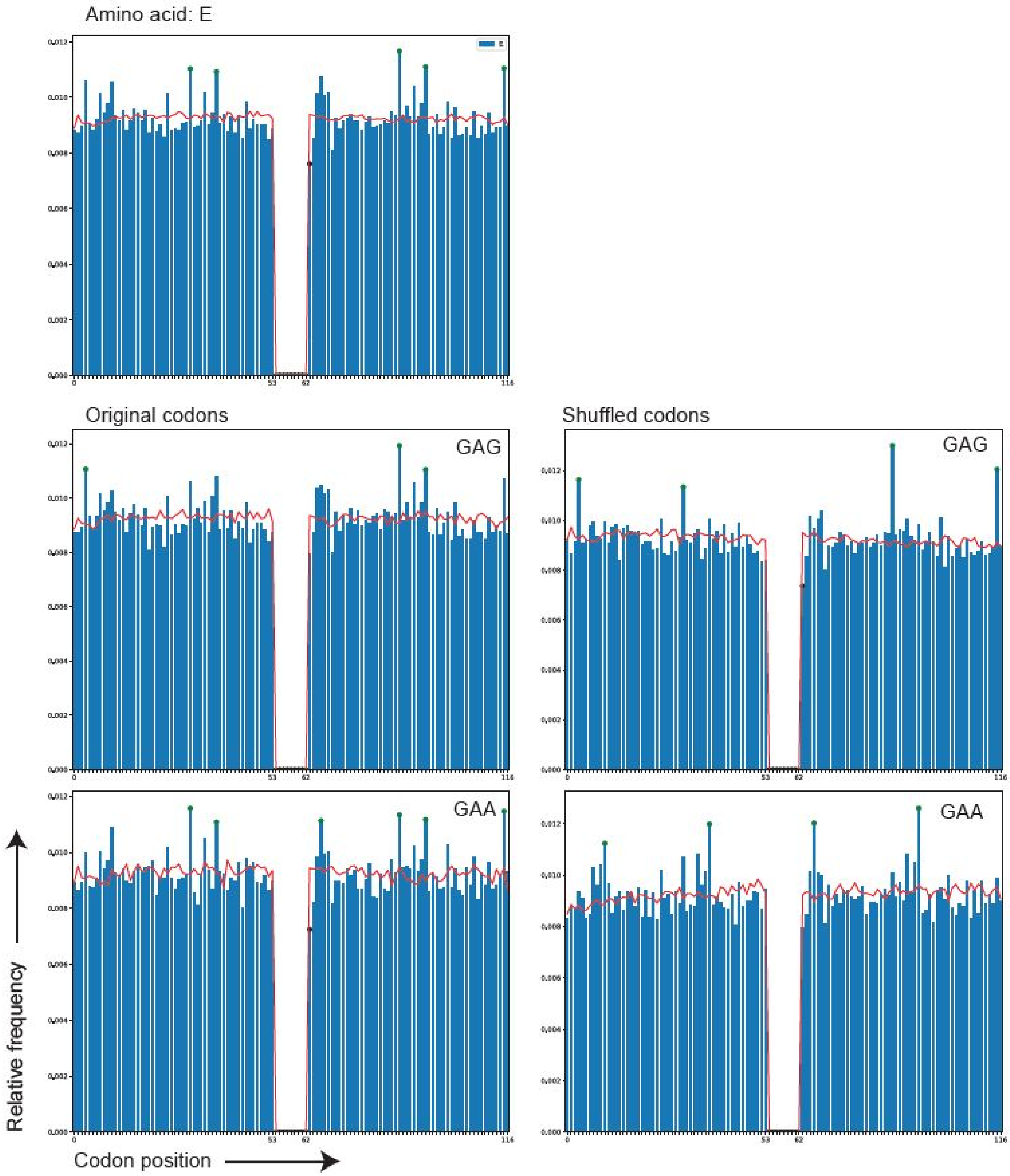


**Supplementary Figure S3. Continued.** (D) Glutamic acid (E).

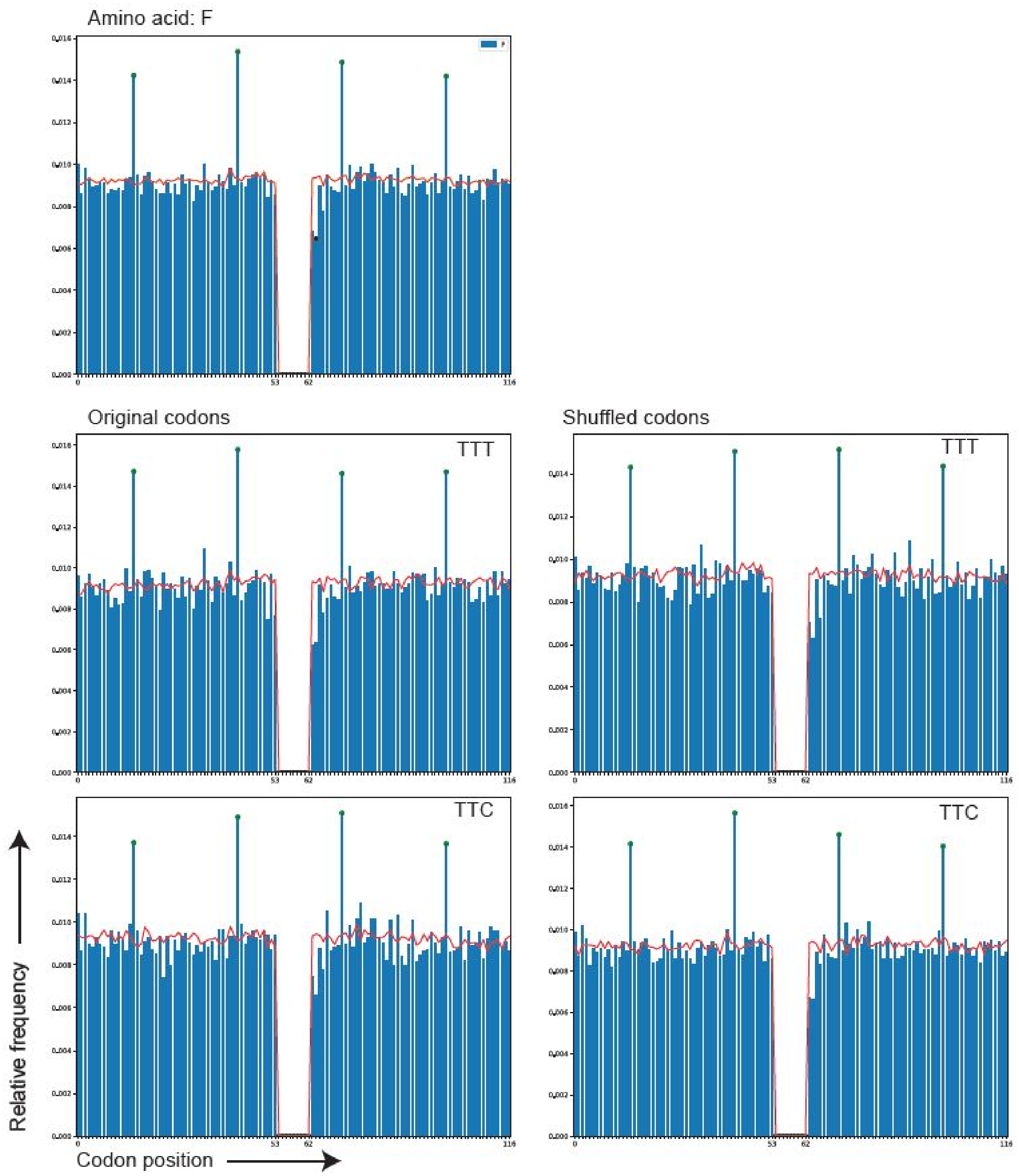


**Supplementary Figure S3. Continued.** (E) Phenylalanine (F).

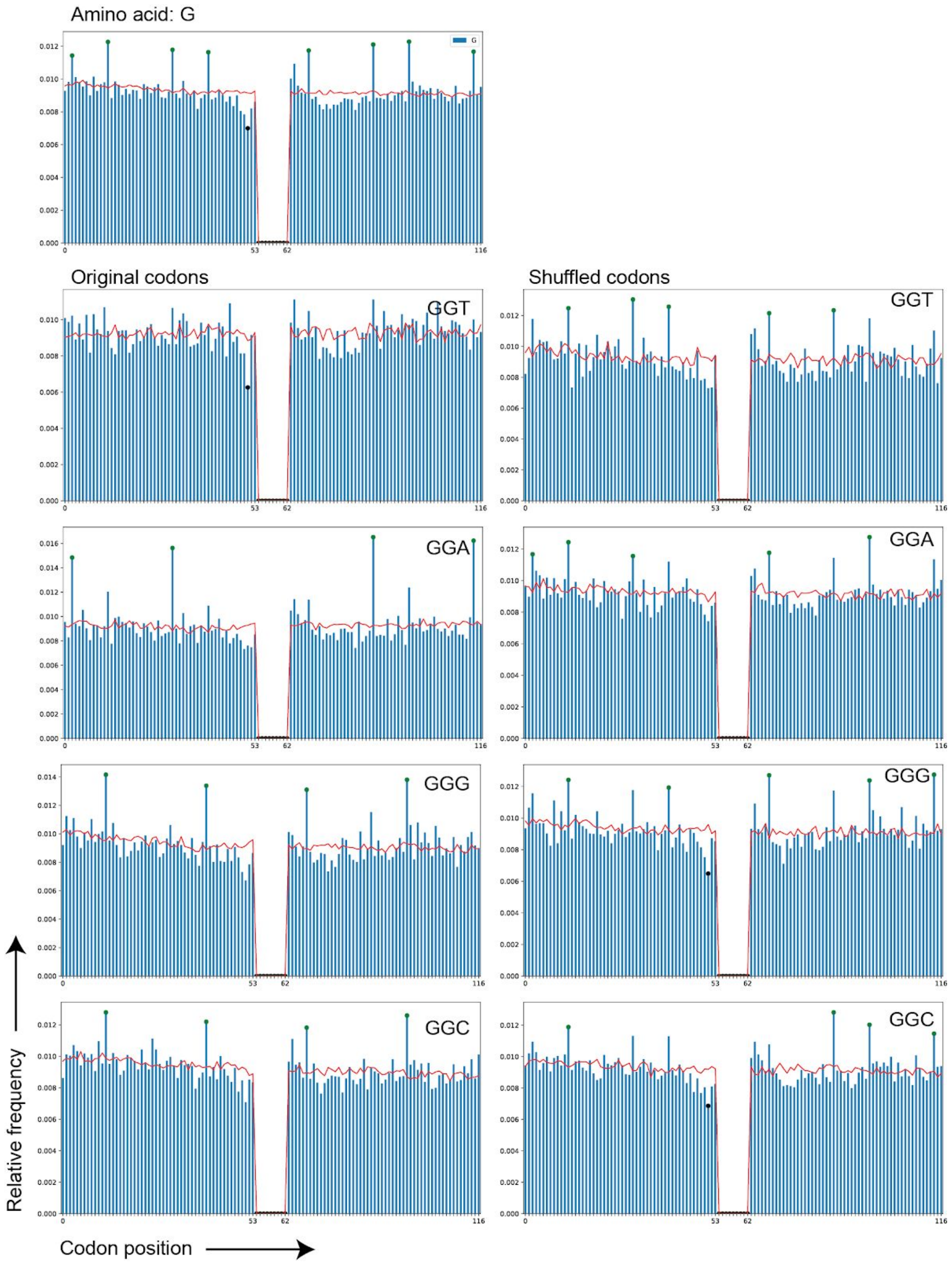


**Supplementary Figure S3. Continued.** (F) Glycine (G).

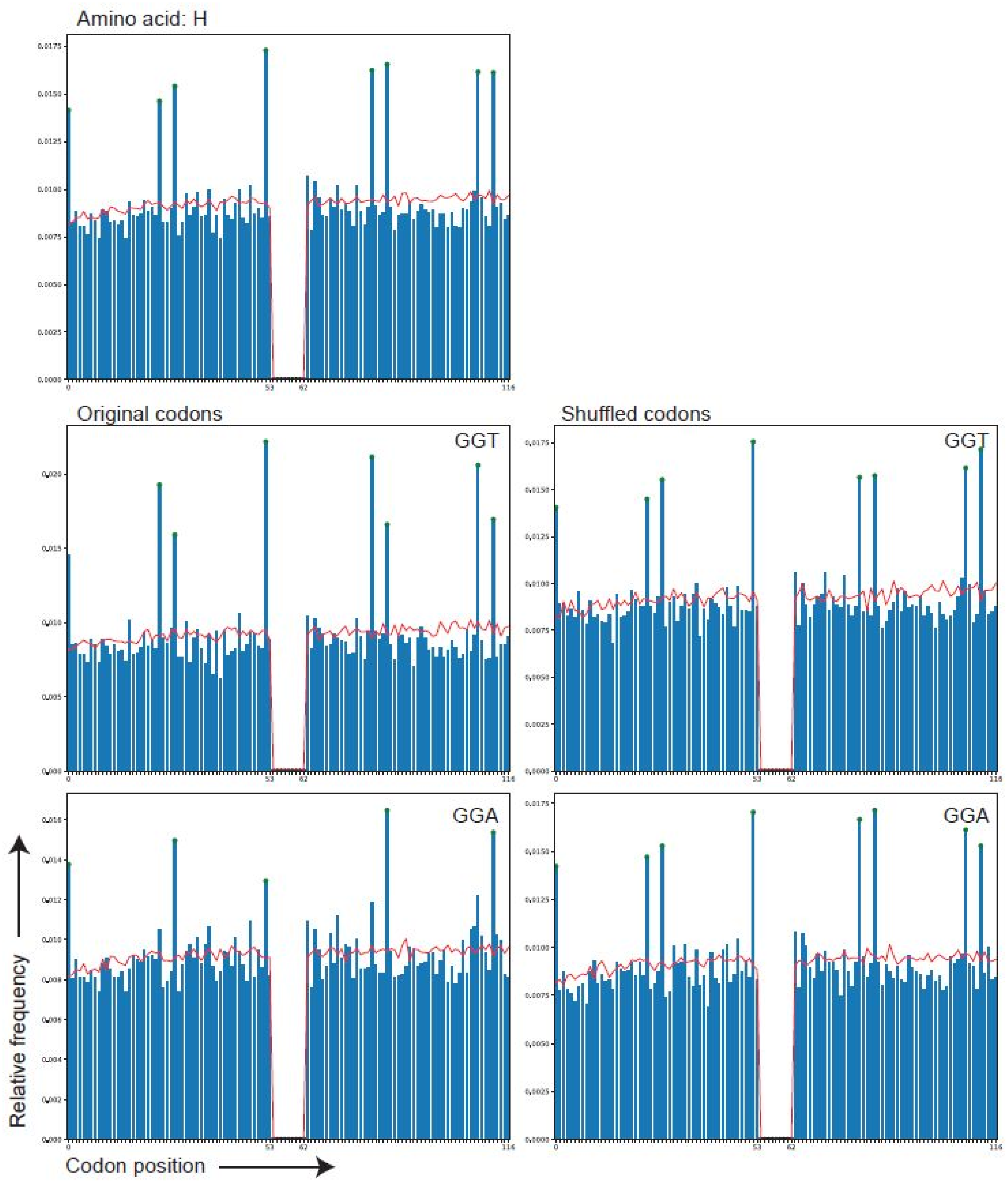


**Supplementary Figure S3. Continued.** (G) Histidine (H).

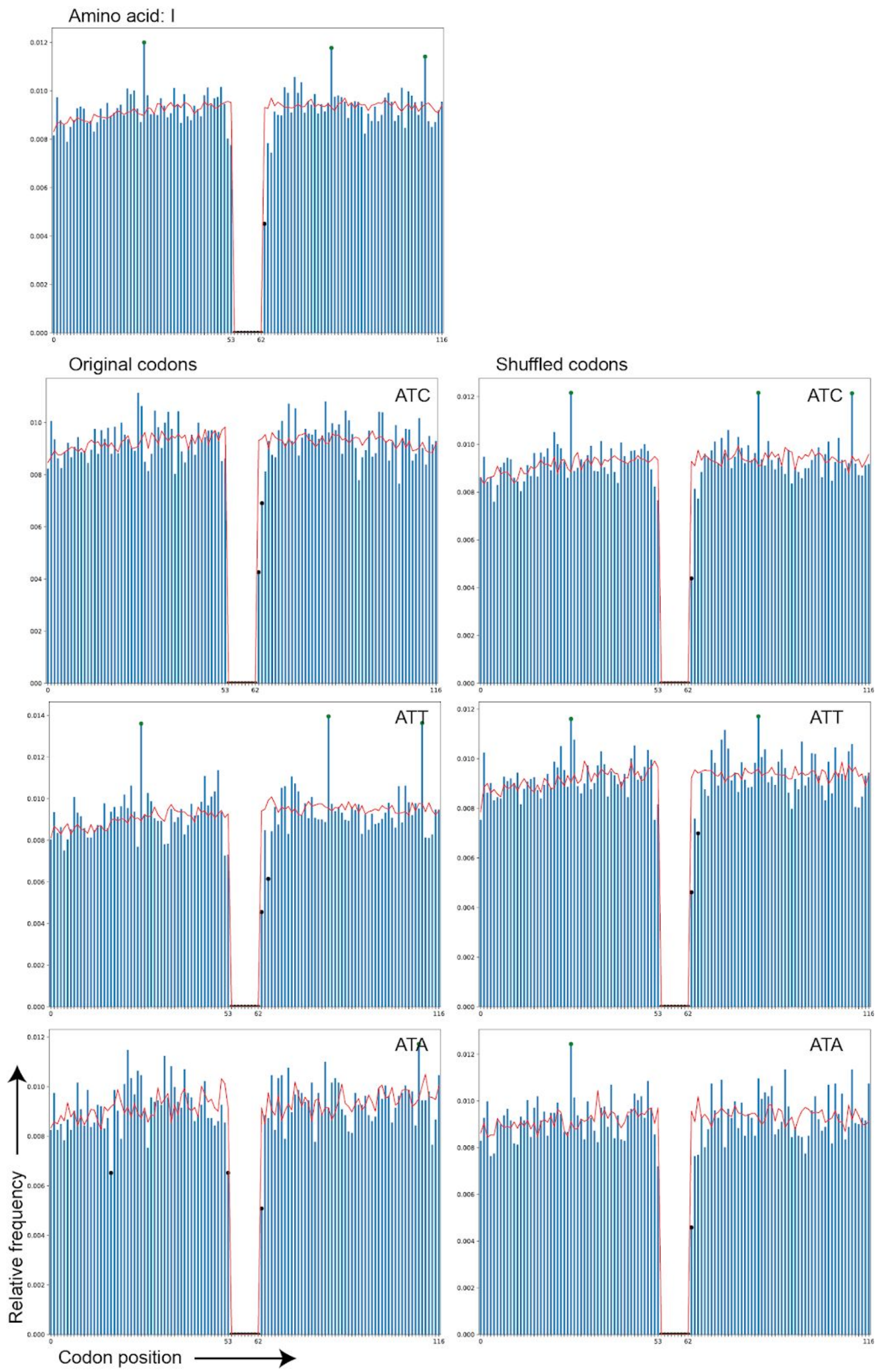


**Supplementary Figure S3. Continued.** (H) Isoleucine (I).

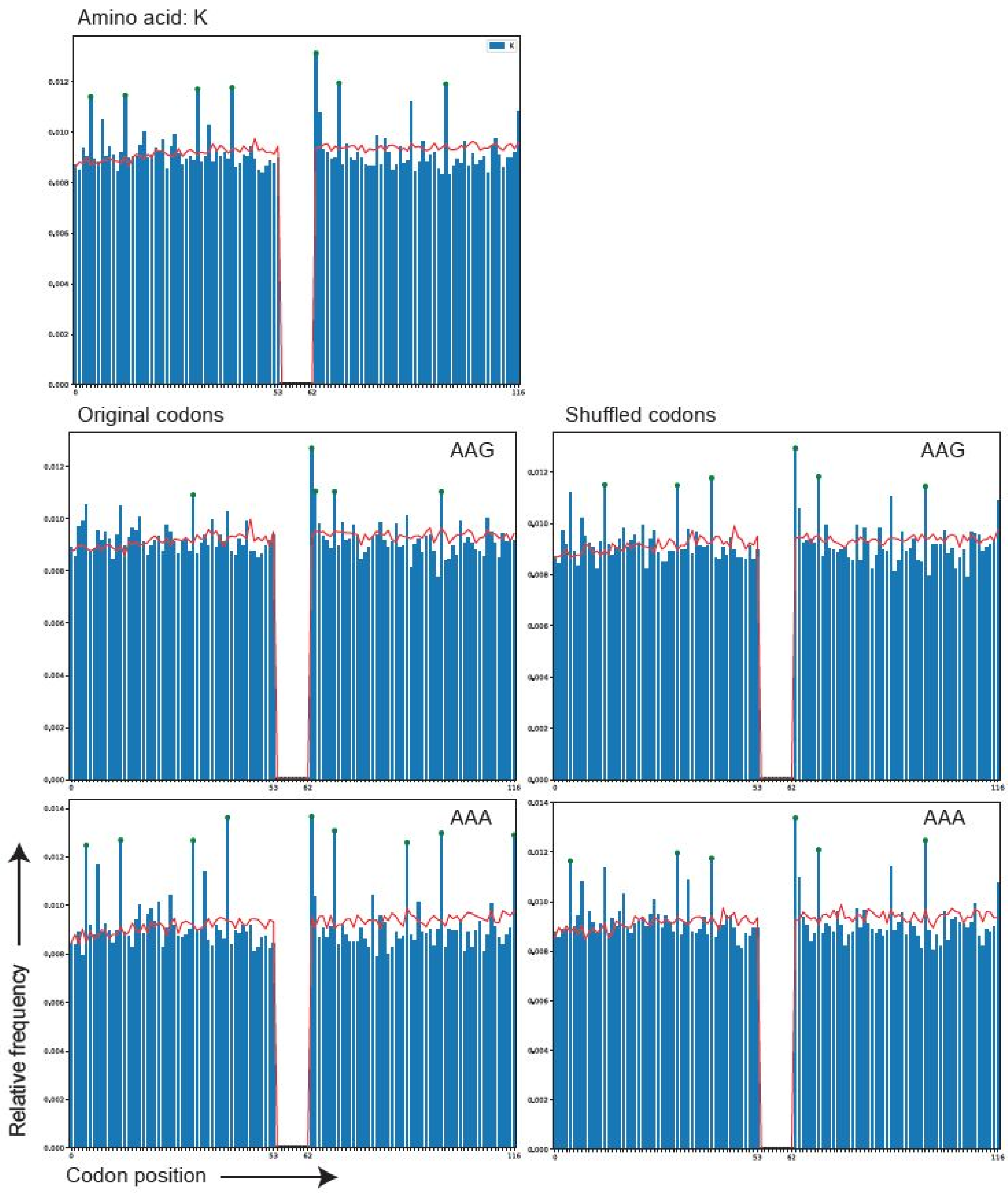


**Supplementary Figure S3. Continued.** (I) Lysine (K).

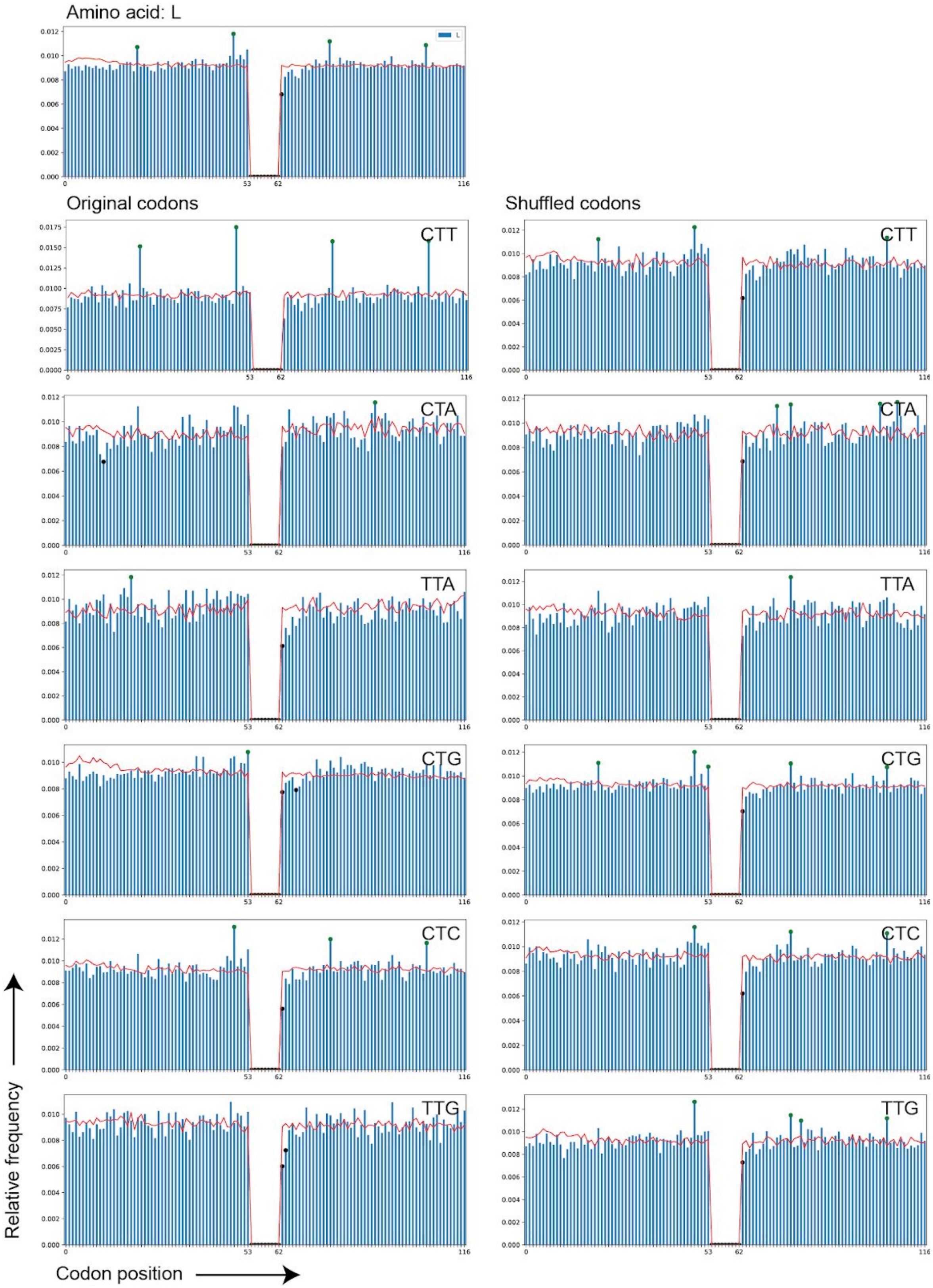


**Supplementary Figure S3. Continued.** (J) Leucine (L).

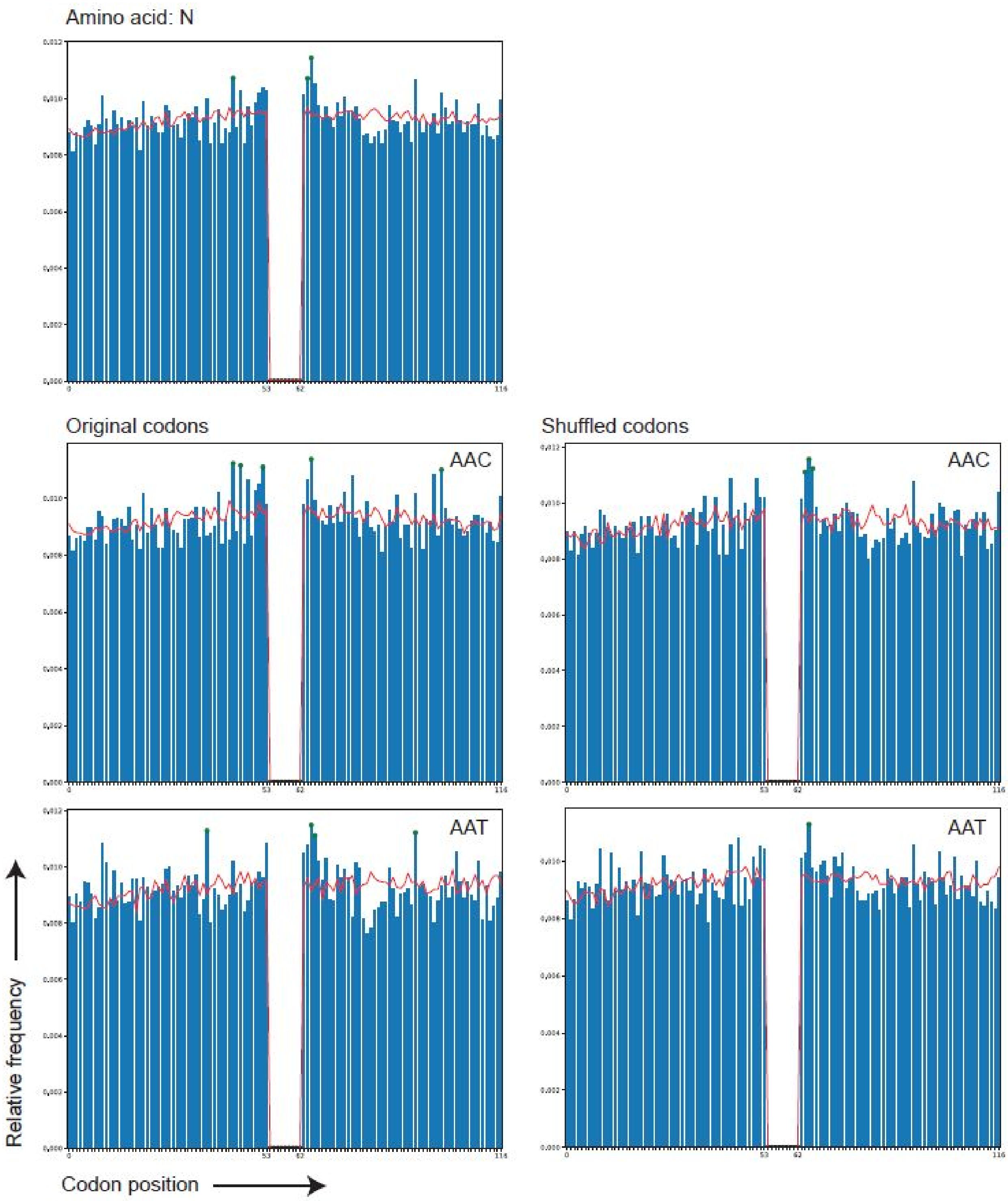


**Supplementary Figure S3. Continued.** (K) Asparagine (N).

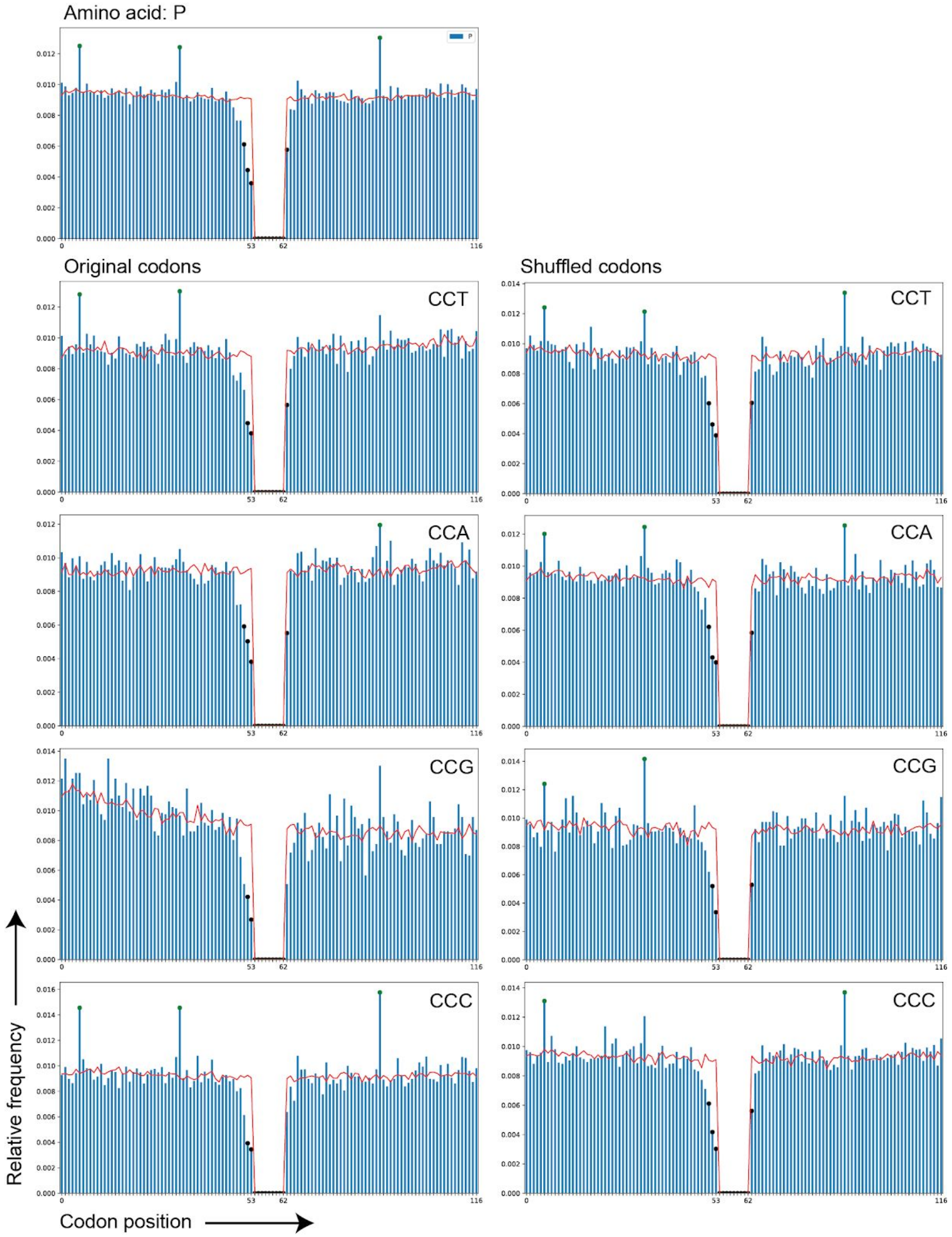


**Supplementary Figure S3. Continued.** (L) Proline (P).

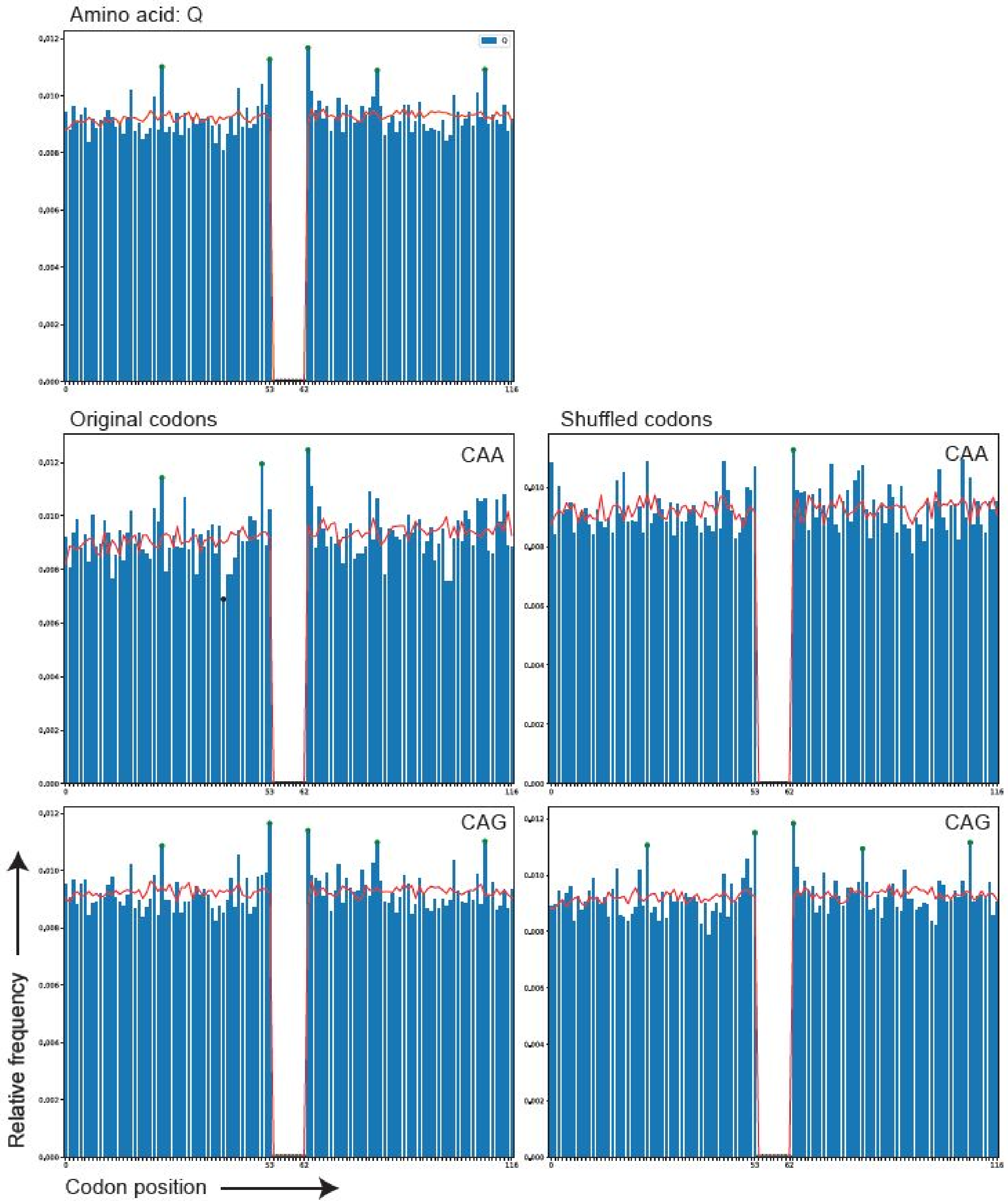


**Supplementary Figure S3. Continued.** (M) Glutamine (Q).

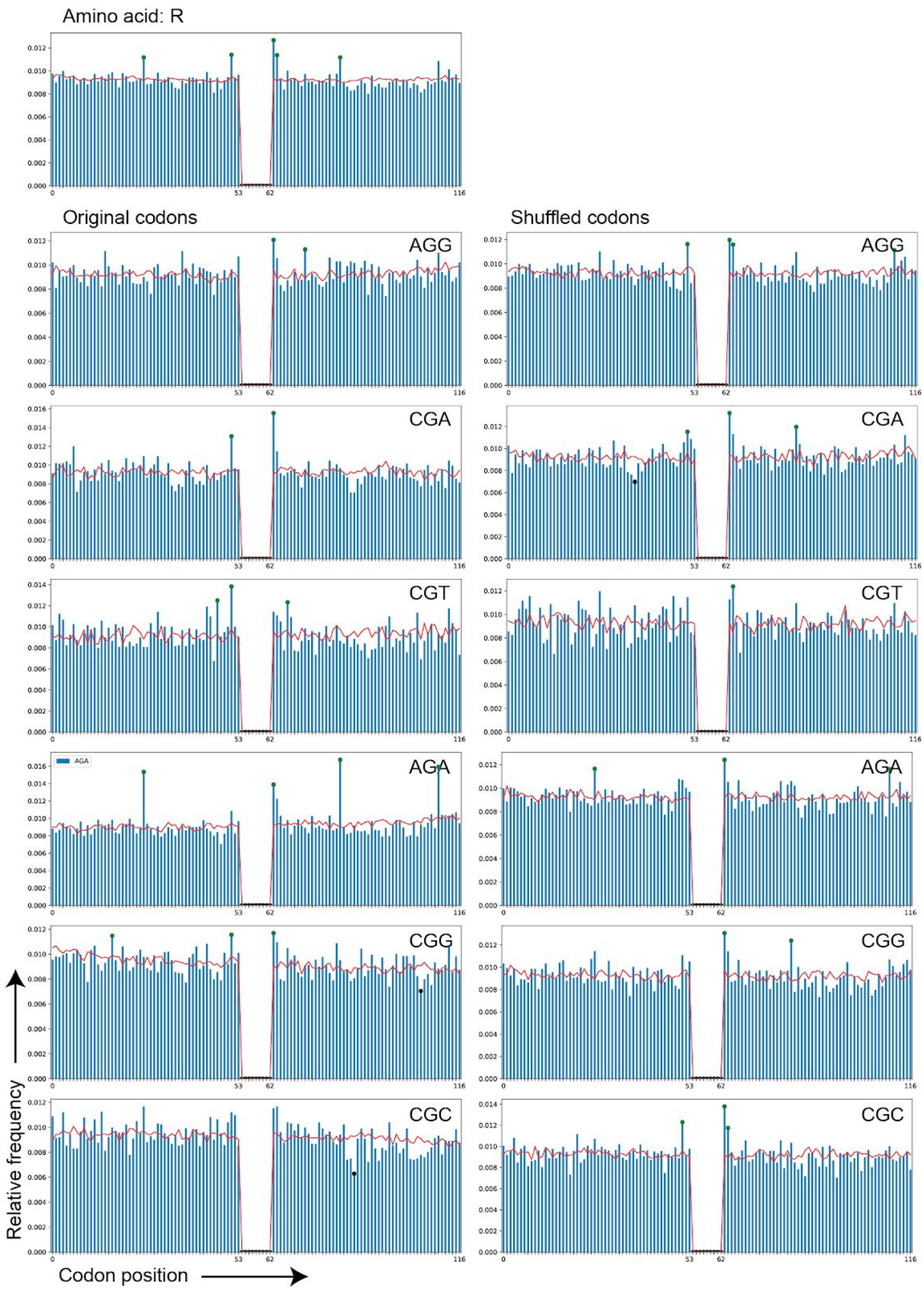


**Supplementary Figure S3. Continued.** (N) Arginine (R).

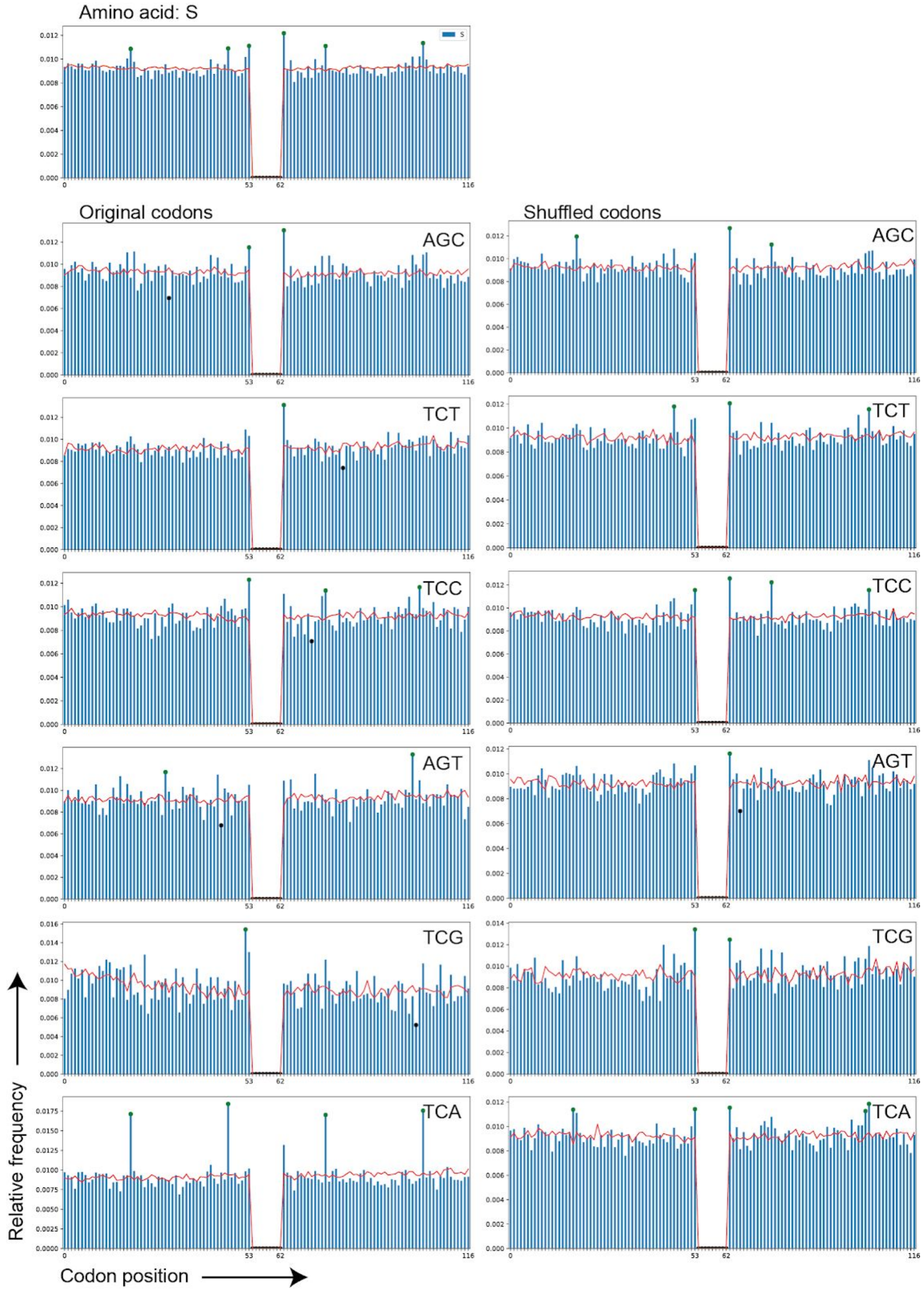


**Supplementary Figure S3. Continued.** (O) Serine (S).

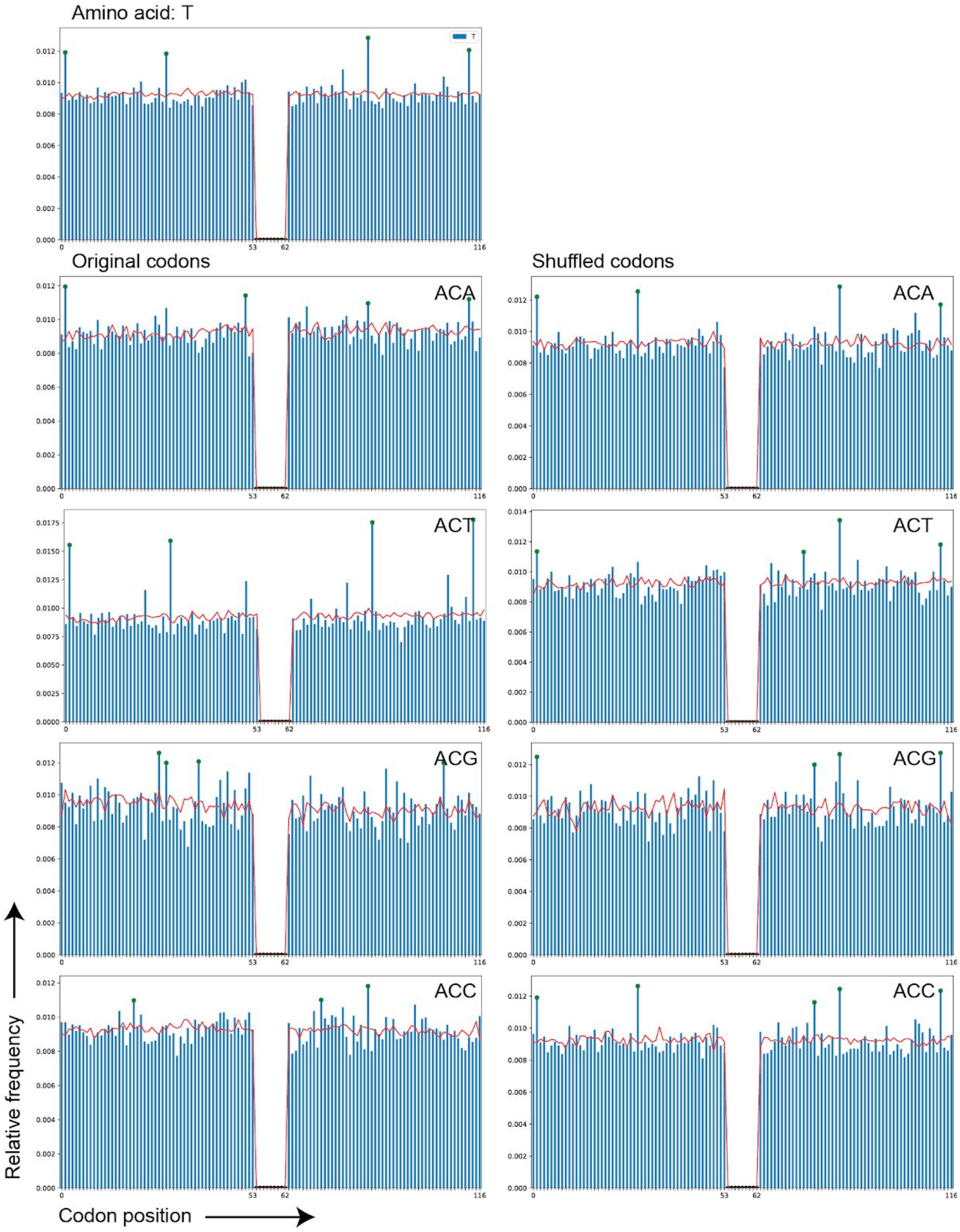


**Supplementary Figure S3. Continued.** (P) Threonine (T).

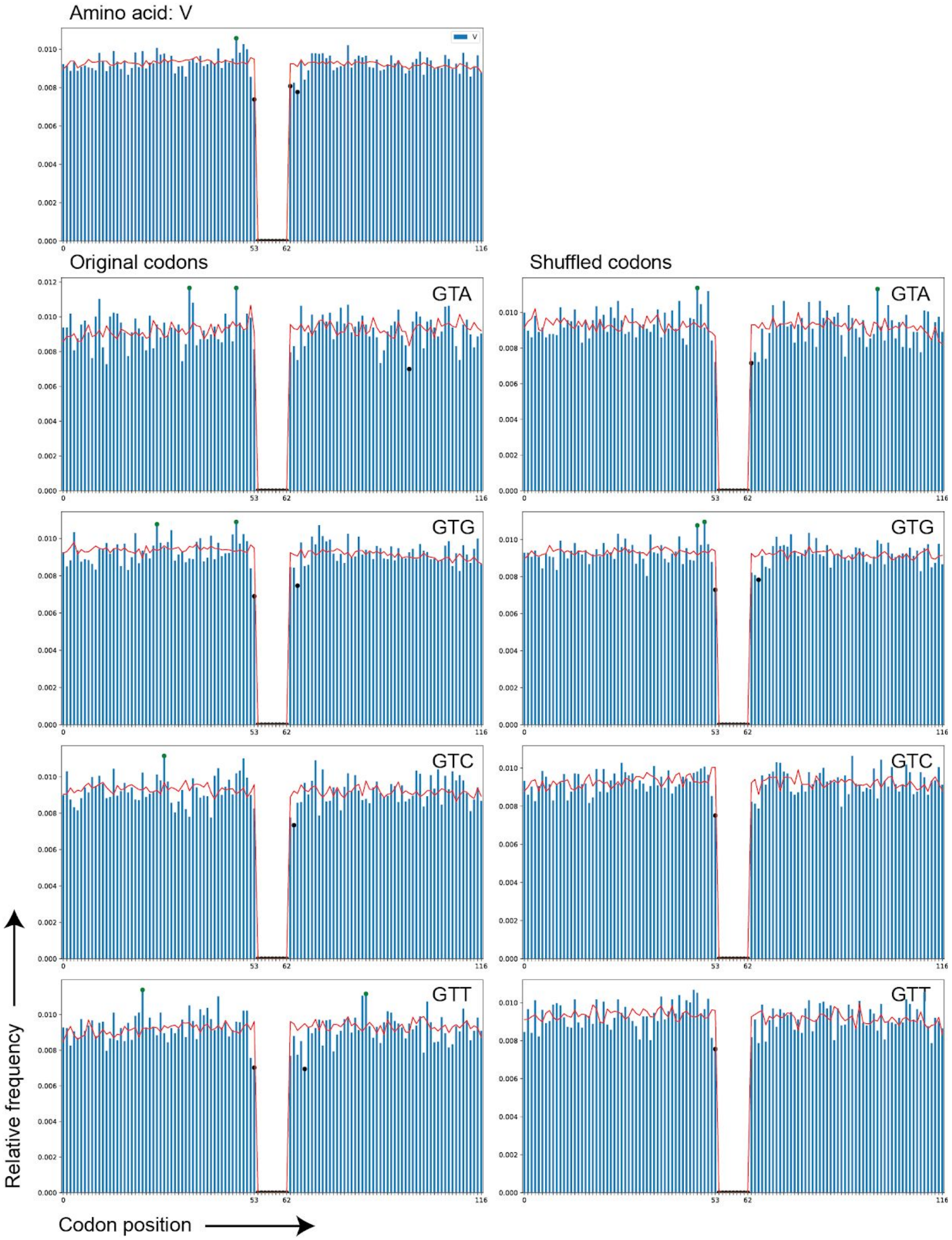


**Supplementary Figure S3. Continued.** (Q) Valine (V).

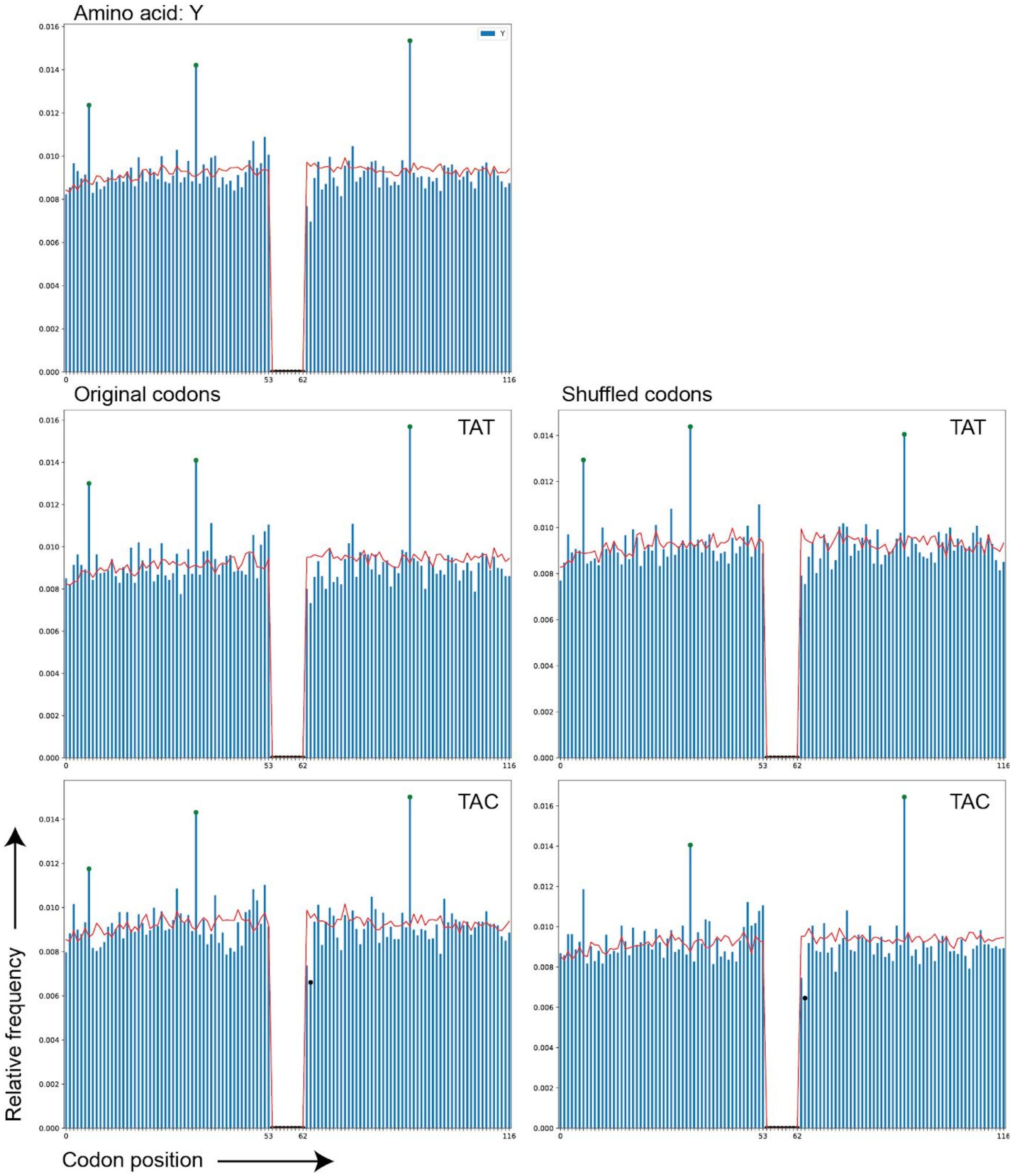


**Supplementary Figure S3. Continued.** (R) Tyrosine (Y).

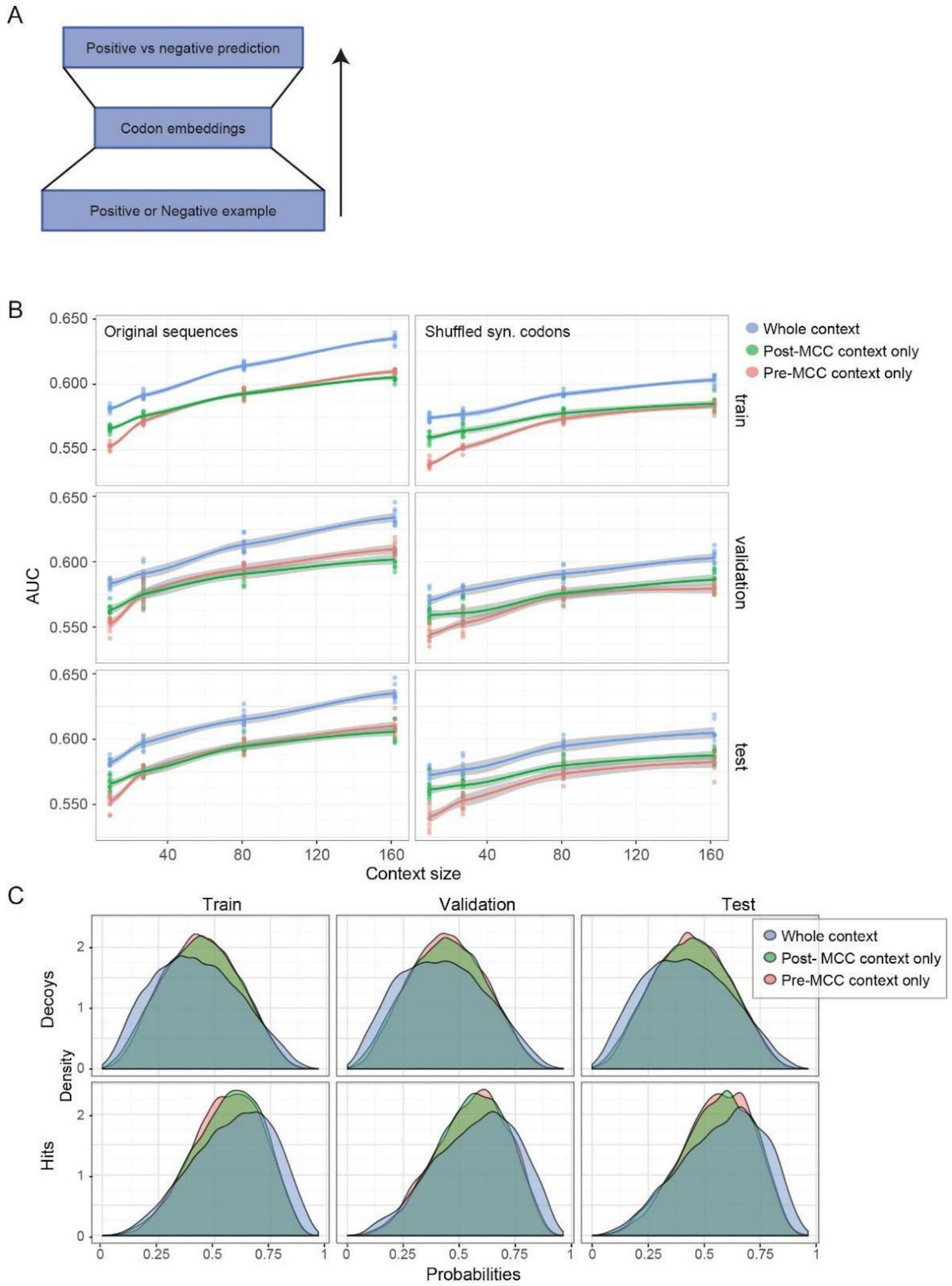


**Supplementary Figure S4.** CAMAP architecture and detailed predictions. (A) Architecture of the ANN used in this work. (B) Results for the AUC on all train, validation and test subsets. Grey areas represent the 95% confidence intervals. (C) Distributions of output probabilities of CAMAPs used to calculate correlations in Supplementary Figure S5.

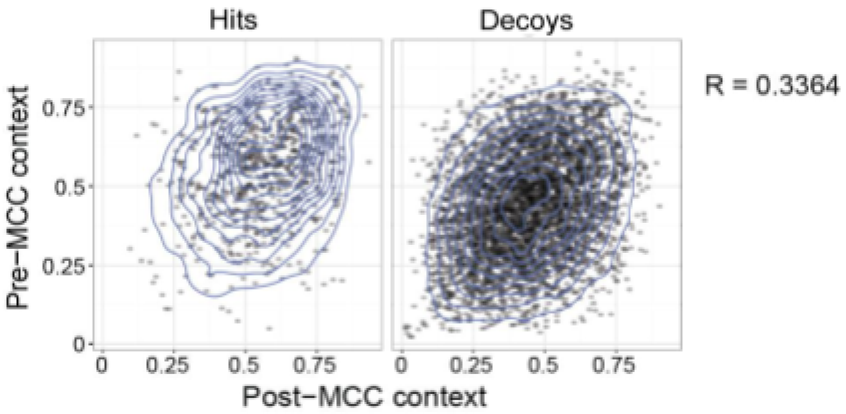


**Supplementary Figure S5**. Correlation between CAMAP prediction score trained only with pre-MCC or post-MCC sequences. For each sequence in the test set we calculated the average prediction score given by CAMAPs in each condition, and calculated the Pearson correlation using the R software. Densities were calculated on all points and drawn using ggplot2. Only a random subset of the points is represented in the figures to limit their size.

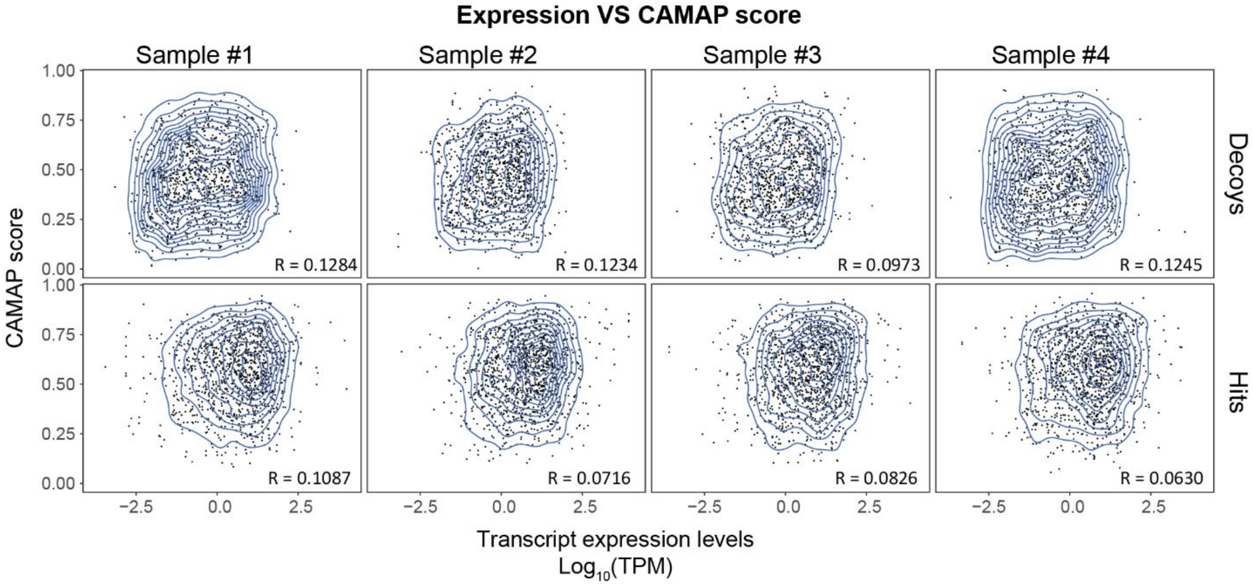


**Supplementary Figure S6.** Absence of correlation between CAMAP prediction score and transcript expression levels in 4 individual B-LCL samples (each derived from a different subject).

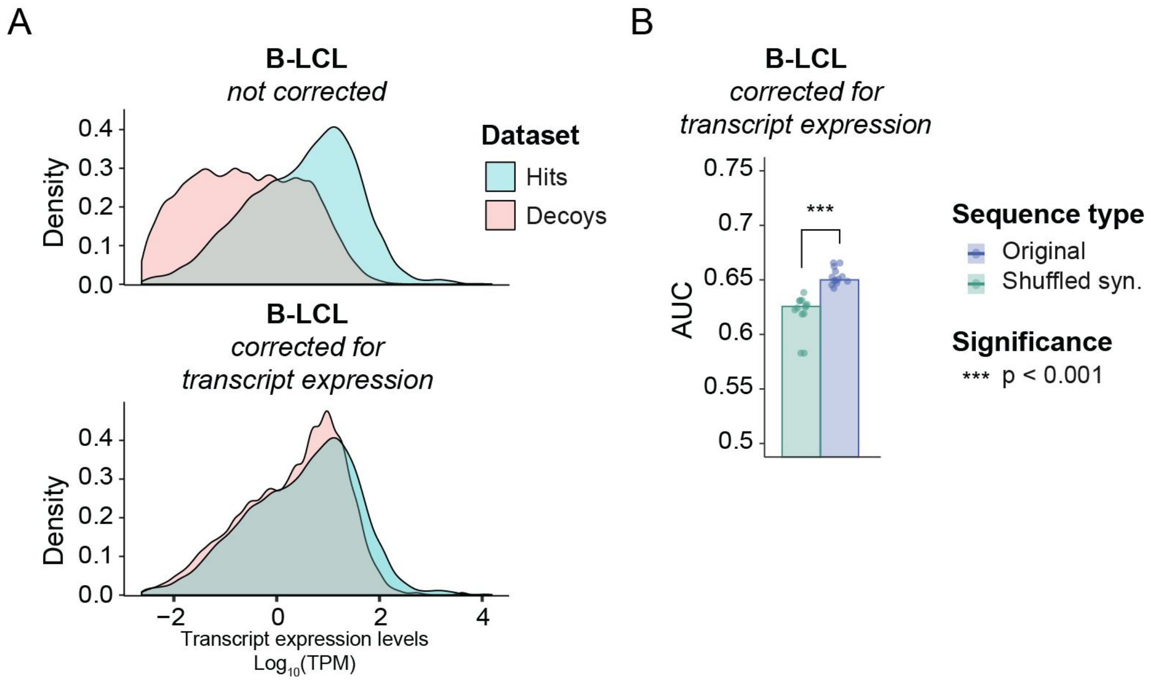


**Supplementary Figure S7. Training of CAMAP on dataset selected to reflect positive dataset's distribution in expression levels.** (A) Distribution of transcript expression levels for normal datasets (related to Figure 2) and the dataset used here to retrain CAMAP. As shown in this figure, the decoy dataset was selected to mirror the distribution of transcript expression level in the hit dataset. (B) CAMAP performance (measured by the AUC) when trained using the decoy dataset that mirrors the transcript expression levels of the hit dataset. Significance was assessed using bilateral paired Student T test ($p = 5.36 \times 10^{-7}$).

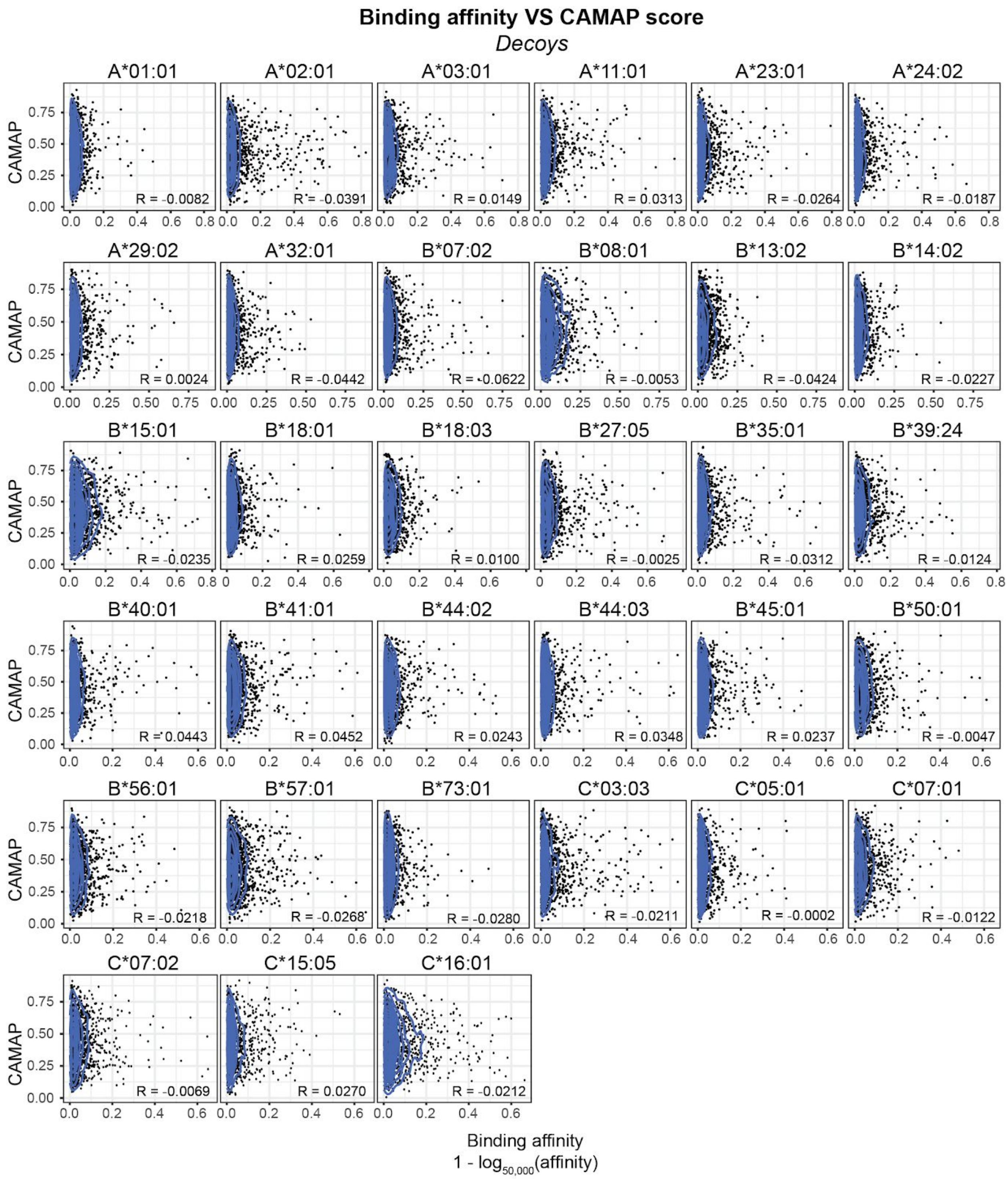


**Supplementary Figure S8A.** Absence of correlation between CAMAP prediction score and binding affinities for individual alleles for decoys.

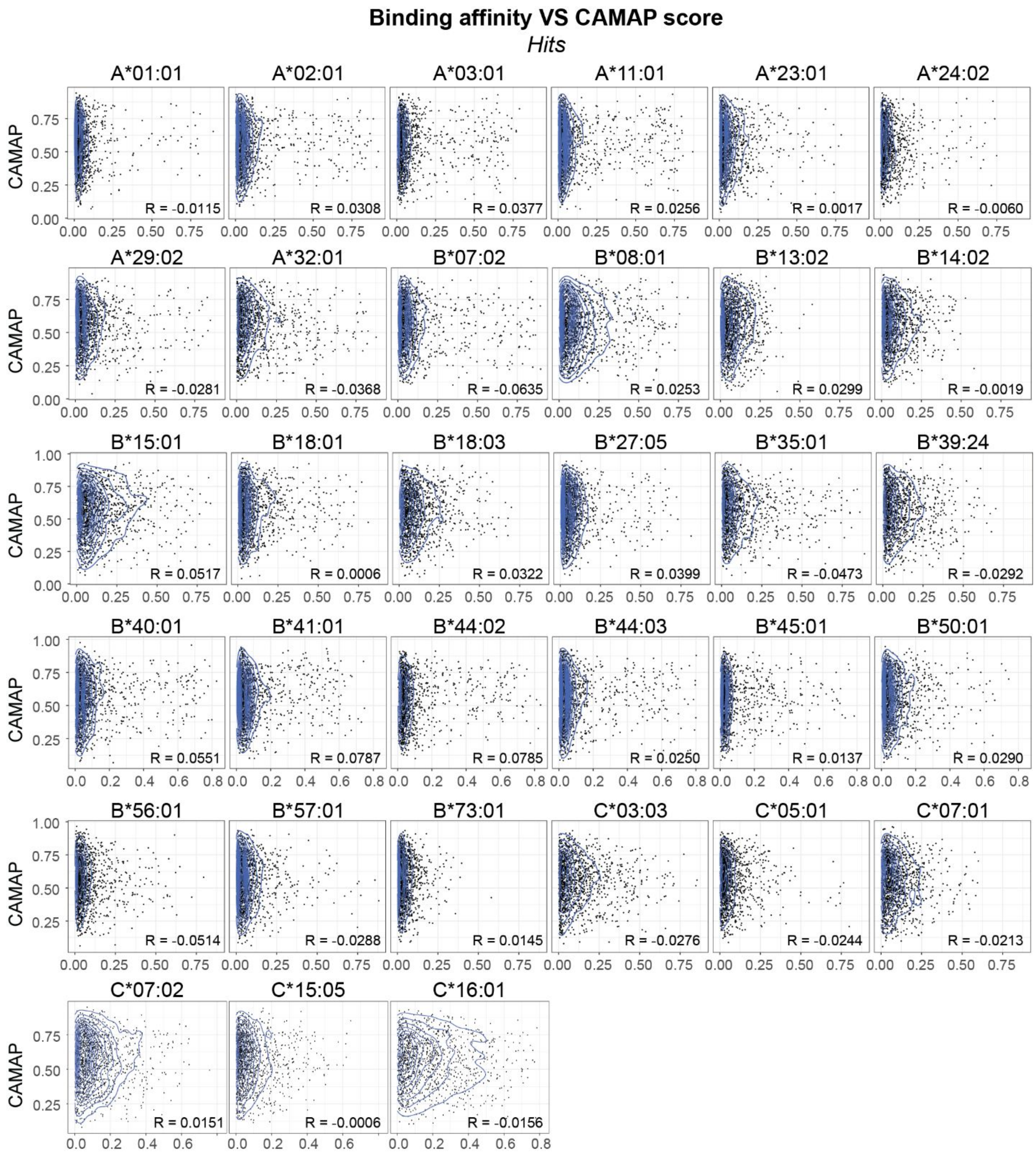


**Supplementary Figure S8B.** Absence of correlation between CAMAP prediction score and binding affinities for individual alleles for hits.

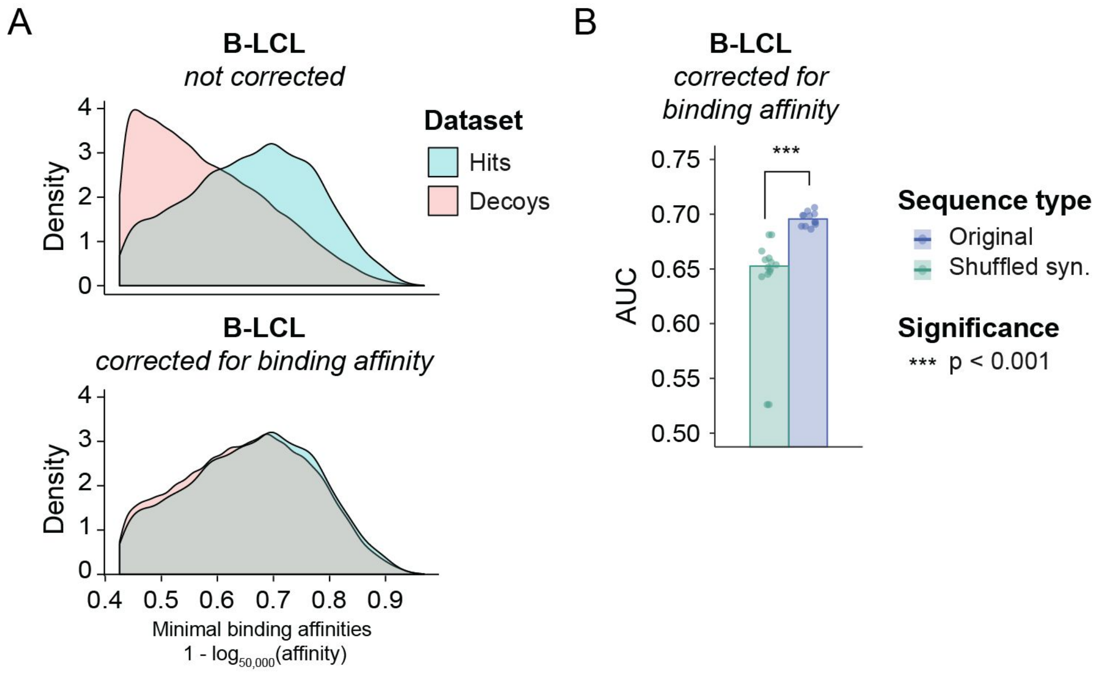


**Supplementary Figure S9. Training of CAMAP on dataset selected to reflect positive dataset's distribution in binding affinities.** (A) Distribution of binding affinities for normal datasets (related to Figure 2) and the corrected dataset used to retrain CAMAP. As shown in this figure, the decoy dataset was selected to mirror the distribution of binding affinities in the hit dataset. (B) CAMAP performance (measured by the AUC) when trained using the decoy dataset that mirrors the binding affinities of the hit dataset. Significance was assessed using bilateral paired Student T test ($p = 1.21 \times 10^{-9}$).

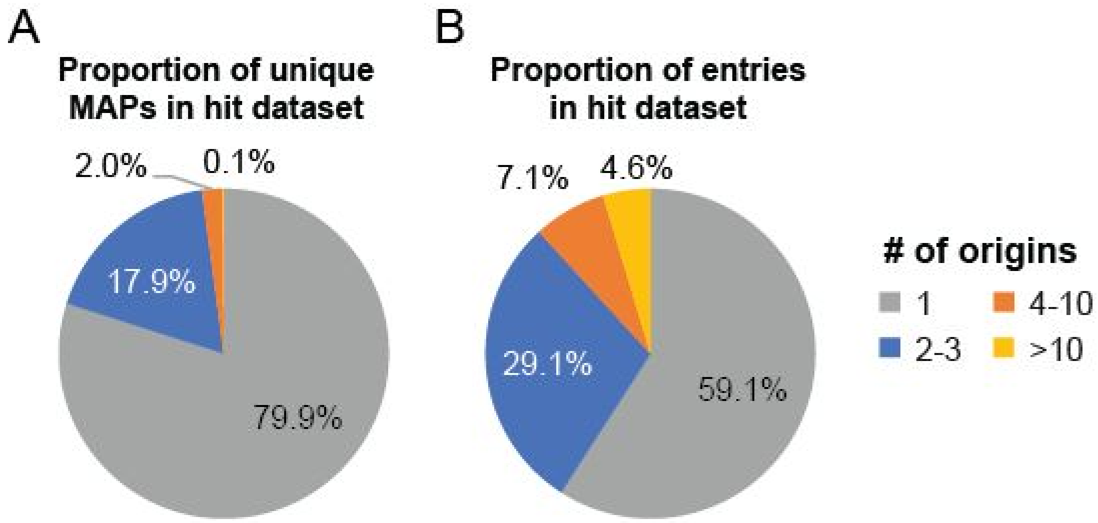


**Supplementary Figure S10. Evaluation of homology in hit dataset and its impact of CAMAP performance.** (A) Proportion of unique MAPs that can be ascribed to a single origin, 2-3, or >10 possible origins. (B) Proportion of entries in the hit dataset that encode for MAPs with a single origin, 2-3, 4-10 or >10 possible origins

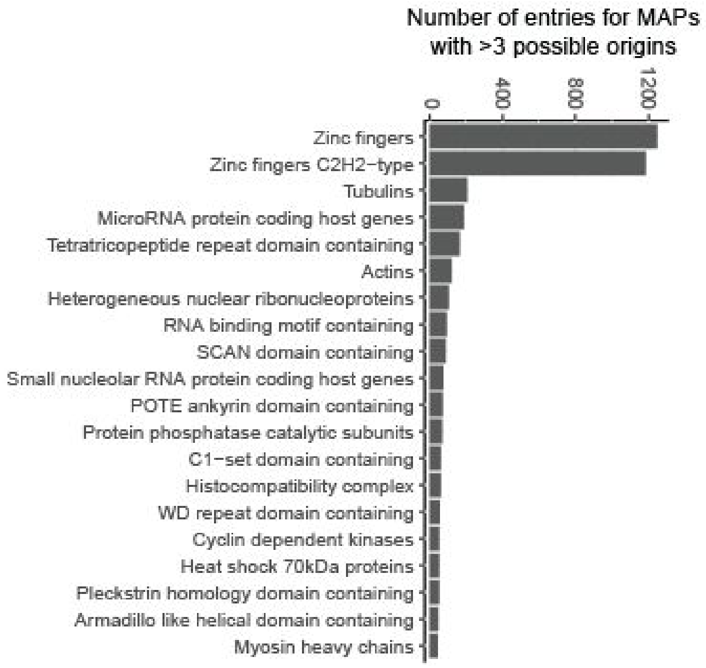


**Supplementary Figure S11. Gene families overrepresented in hits with >3 possible origins.**

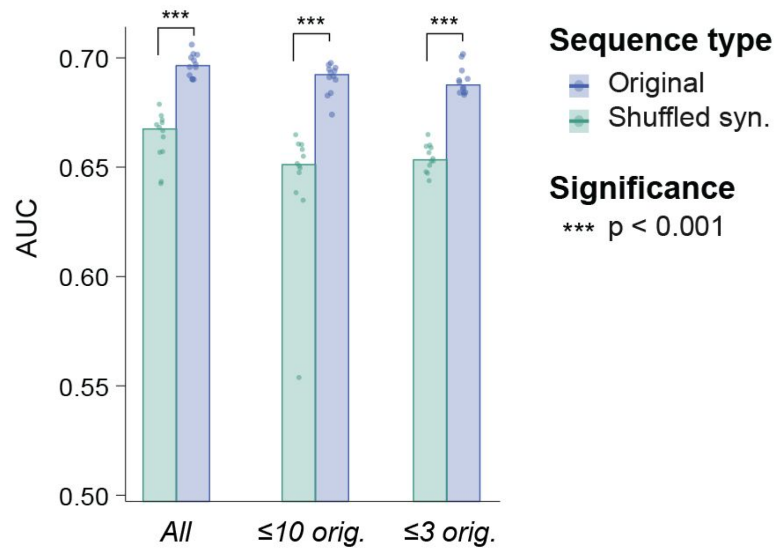


**Supplementary Figure S12. CAMAP performance (AUC) when trained using either all hits (left), hits with 10 possible origins or less (center) or hits with 3 possible origins or less (right).**

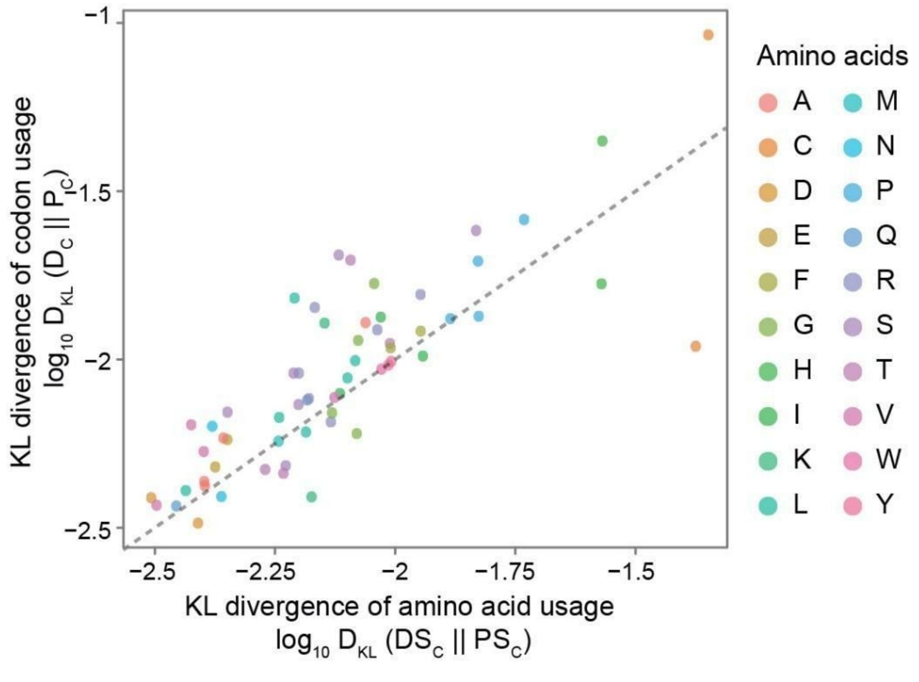


**Supplementary Figure S13**. Kullback-Leibler divergence between hit and decoy datasets in original codon (y-axis) or shuffled synonymous codon sequences (x-axis). Shuffled sequences represent amino acid usage, as codon-specific information are removed with synonymous codon shuffling.

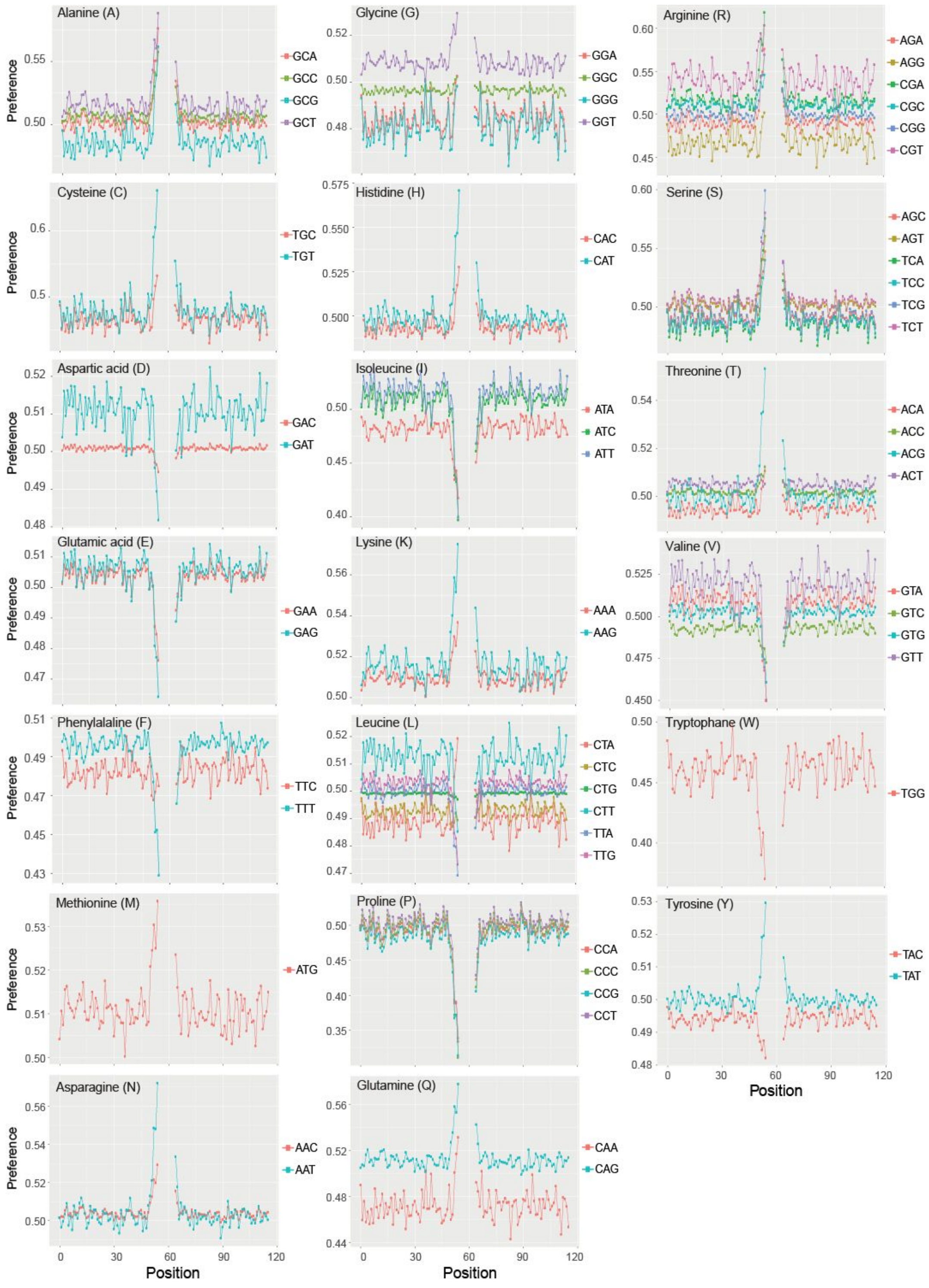

**Supplementary Figure S14.** Preferences per position for all codons for CAMAP trained with original sequences. See Experimental Procedures for more details.

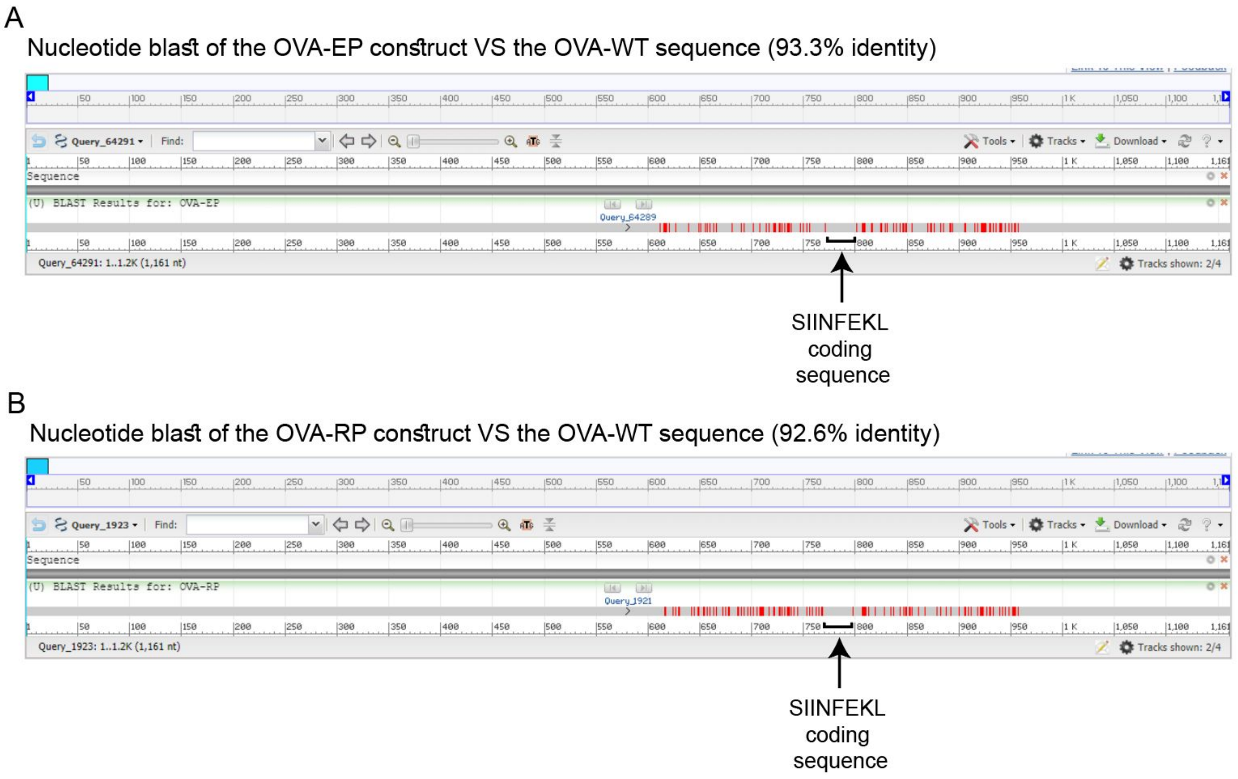


**Supplementary Figure S15.** OVA-construct alignment, showing point mutations (red lines) in the mRNA sequences flanking the SIINFEKL MCC. (A) Comparison of the OVA-EP nucleotide sequence to the wildtype OVA sequence. The OVA-EP and OVA-WT sequences have 93.3% nucleotide identity for a total of 78 modified nucleotides. (B) Comparison of the OVA-RP nucleotide sequence to the wildtype OVA sequence. The OVA-EP and OVA-WT sequences have 92.6% nucleotide identity, for a total of 86 modified nucleotides. Mutations, shown in red, are located only in the 162 nucleotide regions flanking the SIINFEKL coding codons. Of note, the SIINFEKL coding codons (nucleotides 772-799) were not modified between the 3 constructs.

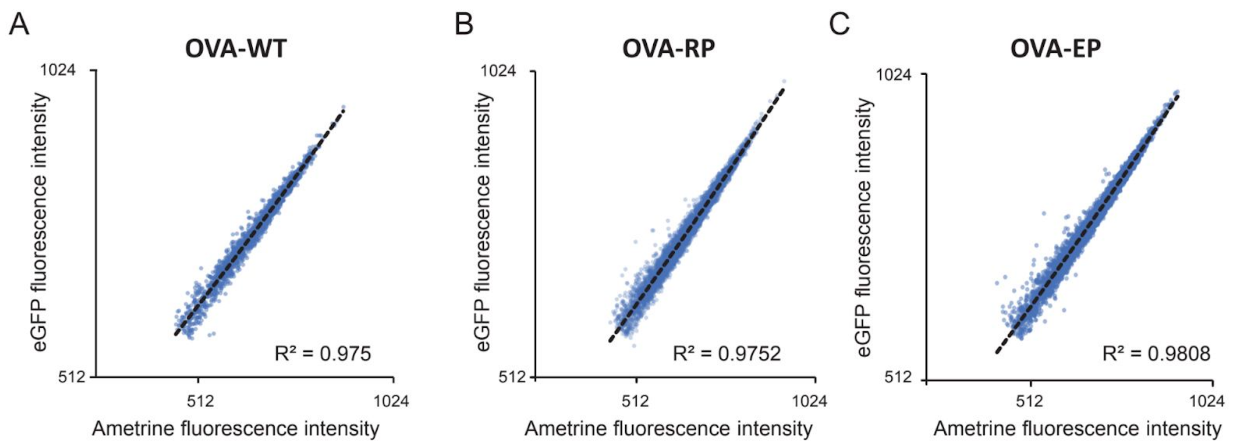


**Supplementary Figure S16.** Correlations between eGFP and Ametrine fluorescence intensity at the single cell level. Single cell eGFP and Ametrine fluorescence intensities measured at 10 hours post-induction are shown for the OVA-WT (A), OVA-EP (B) and OVA-RP (C) constructs. N.B.: only transduced cells are shown (eGFP+ cells).

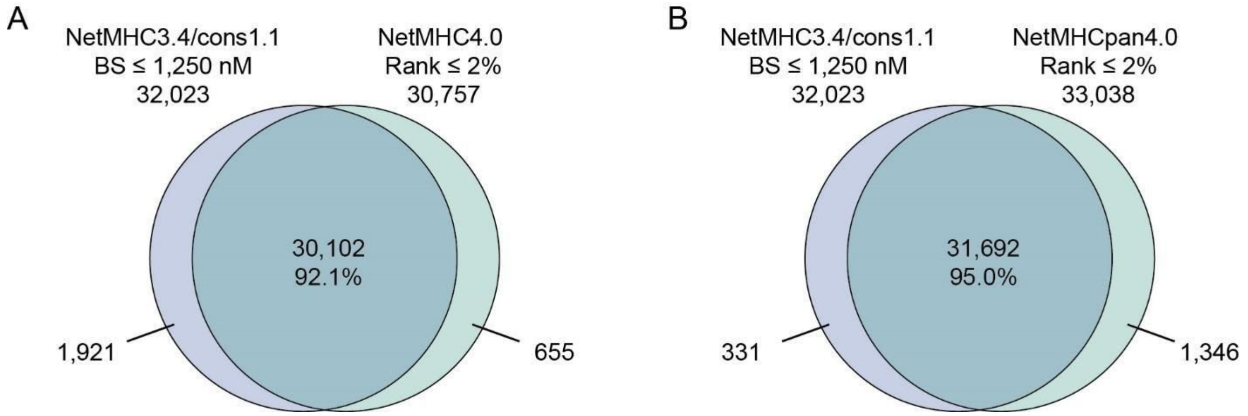


**Supplementary Figure S17.** Validation of MHC-I associated peptides (MAP) dataset from Pearson H. *et al*. (2016) using the new versions of MAP binding affinity prediction algorithm NetMHC4.0 (A) and NetMHCpan4.0 (B).

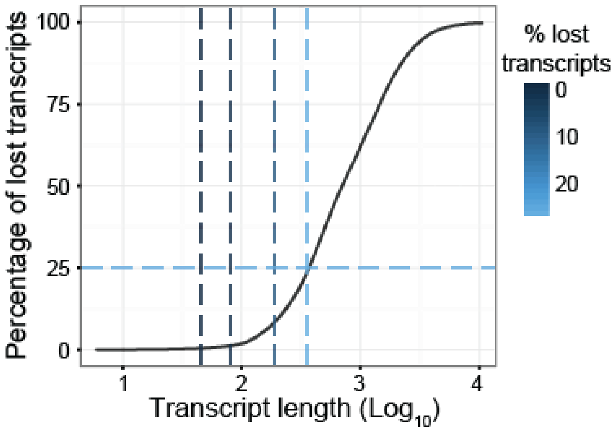


**Supplementary Figure S18.** Percentage of transcript ineligibility as a function of context size. Transcript length corresponds to *C x 2 + 27*, where *C* is the context size in nucleotides and 27 the length of the MCCs. Related to Figure 1A.

### Supplementary Tables

**Supplementary Table S1. Nucleotide sequences of the EP and RP constructs**. SIINFEKL MCC is shown in bold, while the variant regions (pre- and post-MCC contexts of 162-nucleotides) are in blue and italics. Related to Fig. 7.

| **OVA-EP** |
| --- |
| ATGGGCTCCATCGGTGCAGCAAGCATGGAATTTTGTTTTGATGTATTCAAGGAGCTCAAAG<br>TCCACCATGCCAATGAGAACATCTTCTACTGCCCCATTGCCATCATGTCAGCTCTAGCCAT<br>GGTATACCTGGGTGCAAAAGACAGCACCAGGACACAAATAAATAAGGTTGTTCGCTTTGA<br>TAAACTTCCAGGATTCGGAGACAGTATTGAAGCTCAGTGTGGCACATCTGTAAACGTTCA<br>CTCTTCACTTAGAGACATCCTCAACCAAATCACCAAACCAAATGATGTTTATTCGTTCAGC<br>CTTGCCAGTAGACTTTATGCTGAAGAGAGATACCCAATCCTGCCAGAATACTTGCAGTGTG<br>TGAAGGAACTGTATAGAGGAGGCTTGGAACCTATCAACTTTCAAACAGCTGCAGATCAAG<br>CCAGAGAGCTCATCAATTCCTGGGTAGAAAGTCAGACAAATGGAATTATCAGAAATGTCC<br>TTCAGCCAAGCTCCGTGGATTCTCAAACTGCAATGGTTCTGGTTAATGCCATTGTCTTCAA<br>AGGACTGTGGGAGAAAGCATTTAAGGATGAAGACACACAAGCAATGCCTTTCAGAGTGA<br>CTGAG*CAGGAGTCTAAGCCTGTTCAGATGATGTATCAGATTGGTCTTTTTCGTGTTGCTTCTATG*<br>*GCTTCTGAGAAGATGAAGATTCTTGAGCTTCCTTTTGCTAGTGGTACTATGTCTATGCTTGTTCTT*<br>*CTTCCTGATGAGGTTTCTGGTCTTGAGCAGCTTGAA***AGTATAATCAACTTTGAAAAACTG***AC*<br>*TGAGTGGACTTCTTCTAACGTTATGGAGGAGCGTAAGATTAAGGTTTATCTTCCTCGTATGAAGAT*<br>*GGAGGAGAAGTATAACCTTACTTCTGTTCTTATGGCTATGGGAATTACTGATGTTTTTCTAGTTC*<br>*TGCTAACCTTAGTGGTATTTCTTCGGCT*GAGAGCCTGAAGATATCTCAAGCTGTCCATGCAGC<br>ACATGCAGAAATCAATGAAGCAGGCAGAGAGGTGGTAGGGTCAGCAGAGGCTGGAGTGG<br>ATGCTGCAAGCGTCTCTGAAGAATTTAGGGCTGACCATCCATTCCTCTTCTGTATCAAGCA<br>CATCGCAACCAACGCCGTTCTCTTCTTTGGCAGATGTGTTTCCCCTTAA |
| **OVA-RP** |
| ATGGGCTCCATCGGTGCAGCAAGCATGGAATTTTGTTTTGATGTATTCAAGGAGCTCAAAG<br>TCCACCATGCCAATGAGAACATCTTCTACTGCCCCATTGCCATCATGTCAGCTCTAGCCAT<br>GGTATACCTGGGTGCAAAAGACAGCACCAGGACACAAATAAATAAGGTTGTTCGCTTTGA<br>TAAACTTCCAGGATTCGGAGACAGTATTGAAGCTCAGTGTGGCACATCTGTAAACGTTCA<br>CTCTTCACTTAGAGACATCCTCAACCAAATCACCAAACCAAATGATGTTTATTCGTTCAGC<br>CTTGCCAGTAGACTTTATGCTGAAGAGAGATACCCAATCCTGCCAGAATACTTGCAGTGTG<br>TGAAGGAACTGTATAGAGGAGGCTTGGAACCTATCAACTTTCAAACAGCTGCAGATCAAG |

CCAGAGAGCTCATCAATTCCTGGGTAGAAAGTCAGACAAATGGAATTATCAGAAATGTCC
TTCAGCCAAGCTCCGTGGATTCTCAAACTGCAATGGTTCTGGTTAATGCCATTGTCTTCAA
AGGACTGTGGGAGAAAGCATTTAAGGATGAAGACACACAAGCAATGCCTTTCAGAGTGA
CTGAG*CAAGAATCCAAACCGGTCCAAATGATGTACCAAATAGGGCTATTCAGGGTCGCGTCCAT*
*GGCGTCCGAAAAAATGAAAATACTAGAACTACCGTTCGCGTCAGGGACGATGTCCATGCTCGTC*
*CTACTACCGGACGAAGTCTCCGGACTCGAACAACTCGAG***AGTATAATCAACTTTGAAAAAC**
**TG***ACAGAATGGACATCCTCCAATGTCATGGAAGAAAGGAAAATAAAAGTCTACCTCCCGAGGAT*
*GAAAATGGAAGAAAAATACAATCTAACATCCGTCCTAATGGCGATGGGTATAACAGACGTCTTCT*
*CCTCATCCGCGAATCTATCAGGGATATCCAGCGCG*GAGAGCCTGAAGATATCTCAAGCTGTC
CATGCAGCACATGCAGAAATCAATGAAGCAGGCAGAGAGGTGGTAGGGTCAGCAGAGGC
TGGAGTGGATGCTGCAAGCGTCTCTGAAGAATTTAGGGCTGACCATCCATTCCTCTTCTGT
ATCAAGCACATCGCAACCAACGCCGTTCTCTTCTTTGGCAGATGTGTTTCCCCTTAA

**Supplementary Table S2. Number of peptides needed to capture 1%, 5 and 10% of epitopes detected by mass spectrometry in B721.221 and PBMC cell lines.** The lower the number of peptides needed to capture the respective number of epitopes, the better the performance of the prediction model. This is also illustrated by the percentage of false identification (false positive rate, FPR) reported here. Peptides were rank-ordered according to regression scores. Of note, only the maximal regression score was kept for peptides with multiple potential origins.

| Cell lines | Model | n | FPR | n | FPR | n | FPR |
|---|---|---|---|---|---|---|---|
| | | 1% hits (n = 46) | | 5% hits (n = 231) | | 10% hits (n = 462) | |
| **B721.221**<br>_# MAPs:_<br>397,939<br>_# hits:_<br>4625<br>_% hits/entries:_<br>1.16 | NetMHCpan4.0 | 208 ± 26 | 77.9% | 1096 ± 38 | 78.9% | 2362 ± 92 | 80.4% |
| | NetMHCpan4.0+ expression | 85 ± 6 | 45.9% | 426 ± 18 | 45.8% | 895 ± 15 | 48.4% |
| | **NetMHCpan4.0 + expression + CAMAP** | **71 ± 4** | **35.2%** | **380 ± 17** | **39.2%** | **804 ± 20** | **42.5%** |
| | | 1% hits (n = 16) | | 5% hits (n = 81) | | 10% hits (n = 162) | |
| **PBMCs**<br>_# MAPs:_<br>411,468<br>_# hits:_<br>1615<br>_% hits/entries:_<br>0.39 | NetMHCpan4.0 | 122 ± 34 | 86.9% | 824 ± 88 | 90.2% | 1903 ± 132 | 91.5% |
| | NetMHCpan4.0+ expression | 116 ± 10 | 86.2% | 360 ± 22 | 77.5% | 806 ± 34 | 79.9% |
| | **NetMHCpan4.0 + expression + CAMAP** | **62 ± 11** | **74.2%** | **257 ± 33** | **68.5%** | **658 ± 49** | **75.4%** |
